# Rapid adaptation of recombining populations on tunable fitness landscapes

**DOI:** 10.1101/2022.11.16.516770

**Authors:** Juan Li, André Amado, Claudia Bank

## Abstract

How does standing genetic variation affect polygenic adaptation in recombining populations? Despite a large body of work in quantitative genetics, epistatic and weak additive fitness effects among simultaneously segregating genetic variants are difficult to capture experimentally or to predict theoretically. In this study, we simulated adaptation on fitness landscapes with tunable ruggedness driven by standing genetic variation in recombining populations. We confirmed that recombination hinders the movement of a population through a rugged fitness landscape. When surveying the effect of epistasis on the fixation of alleles, we found that the combined effects of high ruggedness and high recombination probabilities lead to preferential fixation of alleles that had a high initial frequency. This indicates that positively epistatic alleles escape from being broken down by recombination when they start at high frequency. We further extract direct selection coefficients and pairwise epistasis along the adaptive path. When taking the final fixed genotype as the reference genetic background, we observe that, along the adaptive path, beneficial direct selection appears stronger and pairwise epistasis weaker than in the underlying fitness landscape. Quantitatively, the ratio of epistasis and direct selection is smaller along the adaptive path (*≈* 1) than expected. Thus, adaptation on a rugged fitness landscape may lead to spurious signals of direct selection generated through epistasis. Our study highlights how the interplay of epistasis and recombination constrains the adaptation of a diverse population to a new environment.

## Introduction

The role of epistasis, which is the genetic-background-dependent effect of mutations on fitness, is a conundrum in evolutionary biology. Because of the complexities it causes, epistasis is difficult to infer and characterize, although it undoubtedly exists in natural traits (Fenster et al. 1997; Crow 2010; Campbell et al. 2018). For instance, strong negative epistasis, manifested as hybrid incompatibility, maintains reproductive isolation between species (Fishman and Sweigart 2018). At the same time, the significance of epistasis during polygenic adaptation is a topic of active debate.

In theoretical biology, models of fitness landscapes, which map genotypes or phenotypes to fitness, have been a recurrent resource for probing the effect of epistasis on adaptation (Bank 2022). Theoretical work on the role of epistasis during adaptation on fitness landscapes has often assumed a strong selection weak mutation regime (SSWM, Gillespie 1984). SSWM implies that only one segregating site is under selection at any time. Under this assumption, only currently beneficial mutations (based on the current underlying genetic background) in an adaptive walk can fix. Thus, interference between selected variants and recombination is neglected. SSWM adaptation on fitness landscapes is well studied and allows for sophisticated analytical characterization of the adaptive process (*e*.*g*., Orr 2002; Bank et al. 2016; Vaishnav et al. 2022). Despite the importance and beauty of the SSWM approximation, natural populations carry standing genetic variation that has evolutionary consequences, the study of which requires the relaxation of the SSWM assumption.

Standing genetic variation (SGV) plays a predominant role in local adaptation (Barrett and Schluter 2008; Marques et al. 2019; Bomblies and Peichel 2022), especially in the face of rapid environmental change. In such a scenario, the typically low mutation rates are unlikely to be sufficient to facilitate rapid adaptation by new mutations (Orr and Unckless 2014). In contrast, when segregating polymorphism becomes selected in a new environment, adaptation to a new niche can be achieved quickly. An empirical example comes from a songbird species (*Sinosuthora webbiana*, Lai et al. 2019). Here, shared standing variants were shown to facilitate high-altitude adaptation in two songbird populations. Interestingly, even the private single-nucleotide polymorphism candidates of adaptation in the two populations are likely to have preexisted in the ancestral population, as was inferred under consideration of the demography of the populations.

Epistasis causes linkage disequilibrium (LD) among standing genetic variants; thus, excess LD could empirically indicate epistasis. For instance, LD among non-synonymous variants within a gene was shown to be significantly larger than LD among non-synonymous variants between genes, especially among interacting amino acids, but this pattern was not observed for synonymous variants (Stolyarova et al. 2022). However, LD for epistatically interacting gene clusters is altered by recombination. On the one hand, most new or rare mutations rely on recombination to bring them together. On the other hand, recombination may break up LD between positively epistatic variants. This is because recombination shuffles genotypes, favoring the direct selection of individual alleles rather than their combined epistatic effects (Neher and Shraiman 2009). Due to its second-order effect, epistasis can only amplify individual genotypes to a high frequency when its effect is large enough to compensate for the genotypes’ removal due to recombination breakdown (Neher et al. 2013).

How does SGV affect adaptation on a fitness landscape in the presence of recombination? SGV can be shuffled by recombination creating new genotypes and thus opening up new adaptive opportunities. Only a small proportion of the fitness landscape, namely those genotypes which are composed of the segregating alleles, can be traversed by standing variation in the absence of new mutations. However, on a short time scale, this is likely to be a much larger number of genotypes than *de novo* mutation would create, especially when mutation rates are low, like in most organisms. Empirically, it is challenging to infer how a population has used its SGV to respond to environmental change and how epistasis has contributed to shaping the evolutionary trajectory.

Here, we approach this question computationally using a fitness landscape model with tunable epistasis. We study the short-term evolutionary dynamics of a haploid recombining population that evolves on fitness landscapes with varying degrees of epistasis by means of standing genetic variation. We quantify how the recombination probability affects the time until the population becomes monomorphic and the final population fitness, considering different levels of epistasis in the fitness landscape. Moreover, we characterize the fixed alleles and extract the pairwise epistasis along the adaptive path. We discuss our results in light of the contribution of epistasis to evolutionary trajectories in nature.

## Methods

### Modeling rapid adaptation in haploid recombining populations

We model the short-term evolution of a haploid recombining population in the presence of epistasis (Figure 1). We initialize a population in the genotype space that has evolved under neutrality; thus, we draw the segregating sites in the population from a neutral site frequency spectrum distribution (explained further below). When a sudden environmental change occurs, these previously neutral sites become selected according to a fitness landscape model. Since we are interested in the rapid adaptation process that follows the change in environment, we neglect *de novo* mutations and focus on the evolution of the preexisting segregating diversity. We use a Rough-Mount-Fuji fitness landscape model (RMF model Aita et al. 2000) to define the fitness landscape in which the population is evolving. As the population evolves, alleles become successively fixed due to selection and drift. The adaptation process ends when the population is monomorphic.

**Figure 1.**
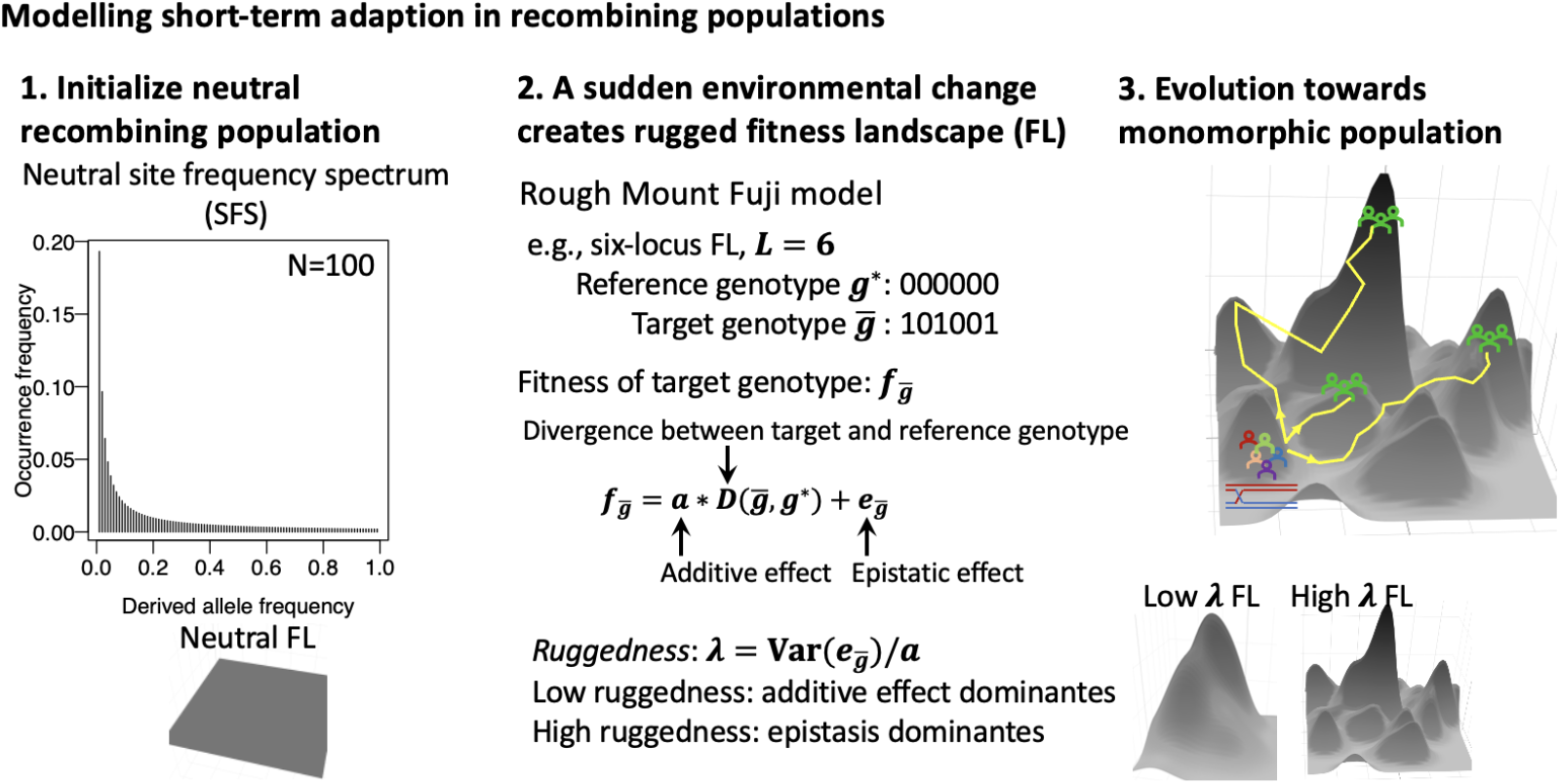
Illustration of the modeling approach. 1. We assume that SGV arose from neutral evolution. 2. Upon environmental change, the population experiences a fitness landscape of the Rough-Mount-Fuji type. 3. Evolution occurs until the population is monomorphic.

In our simulations, each population contains *N* haploid individuals and follows Wright-Fisher dynamics with discrete non-overlapping generations. Every generation, individuals are grouped in pairs and allowed to recombine to form the gamete pool. We assume equidistant loci, such that the recombination probability between two neighboring loci is uniform, and the maximum recombination probability between two loci is 0.5 (here, all loci are independent).

After recombination, *N* individuals are selected from the gamete pool with a probability proportional to their fitness in the fitness landscape. In the main text, we study *N* = 5000 to represent large populations. The supplementary material presents results for smaller populations of *N* = 500 and *N* = 100, representing scenarios of strong(er) genetic drift. For each parameter set, we generated 100 small/low-dimensional and 100 large/higher-dimensional fitness landscapes with 5 and 15 loci, respectively. For each fitness landscape, we simulated adaptation for 100 initial genotype distributions.

### Generating fitness landscapes according to a Rough-Mount-Fuji model

We generate fitness landscapes with a tunable fraction of epistasis using the RMF model (Aita et al. 2000). We assume genotypes with *L* diallelic loci (alleles 0 and 1). Allele 1 at each locus is assigned a constant additive fitness effect *a*. In addition, each genotype receives an epistatic contribution 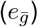 to its fitness. The fitness of genotype 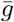 is

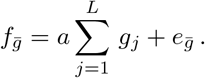

Here, *g*_*j*_ represents the allele at locus *j* (*g*_*j*_ ∈ *{*0, 1*}*). The epistatic term 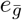 is a random variable drawn from a normal distribution, *N* (0, *σ*^2^), with zero mean and standard deviation *σ*.

After drawing the genotype frequencies of the initial population according to a neutral site frequency spectrum (SFS), as described below, the fitness landscape is parameterized such that minor alleles receive the additive effect *a* = 0.01 (i.e., for each locus, the less frequent allele is assigned the additive beneficial effect). Due to this assignment, the mean fitness of the initial population is by definition lower than the maximum additive fitness. In addition, a random epistatic fitness contribution is assigned to each genotype, drawn from the normal distribution described above. The ratio of the standard deviation of the epistatic effect *σ* to the additive effect *a* determines the ruggedness of the fitness landscape. Low ruggedness implies that the additive effect dominates; high ruggedness indicates that the epistatic effect dominates. We probe fitness landscapes of different ruggedness by studying three values of *σ*, 0.001, 0.01, and 0.1. Hence, the ruggedness of our fitness landscape is 0.1, 1, or 10, which we refer to as smooth, intermediately rugged, and highly rugged fitness landscapes throughout the paper.

### Defining the starting occupancy of the population on the genotype space

Choosing a starting occupancy of the genotype space is nontrivial since the starting frequencies greatly affect the outcome of the dynamics. To create a realistic initial frequency distribution of alleles and genotypes in the genotype space, we assume that previous neutral evolution has generated a neutral SFS from which we draw allele frequencies for each locus, which are combined into genotype frequencies under the assumption of linkage equilibrium. Moreover, all alleles are defined to have an initial frequency above a genetic drift threshold, as detailed in the following subsection. This assumption reduces the homogenizing effect of random genetic drift in the first generations of our simulations. In the following, we describe these assumptions in detail.

#### Site frequency spectrum in the initial population

We consider a haploid population with *N* individuals, originally at neutral mutation-drift equilibrium, suddenly exposed to a fitness landscape. To initialize the population, we adopt the folded neutral SFS in a discrete form (Hudson 2015). We define a minor and major allele for each locus, corresponding to the allele with the smaller and larger frequency in the population, respectively. In a population with size *N*, minor allele frequencies are 1*/N*, 2*/N*, …, *N/*2*N* (when *N* is even, or (*N* − 1)*/*2*N* if *N* is odd). Below, we assume *N* is an even number. The relative abundance of an allele at frequency *i* is 1*/i* + 1*/*(*N* − *i*) (Hudson 2015). Thus, alleles at lower frequencies are more abundant. The probability that a given allele appears *i* times in the population is given by 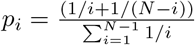, summing the terms corresponding to *i* and *N* − *I* (because the SFS is folded) and normalizing the distribution. Therefore, the mean of minor allele frequency *i* is 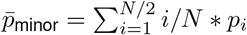, and 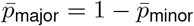. Both *p*_major_ and *p*_minor_ are arithmetic means. The variance of the minor allele frequencies is 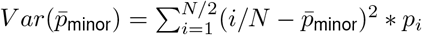, so the standard deviation is the square root of its variance. The mean and variance decrease with increasing population size (Figure S1). The lowest allele frequency, 1*/N*, shifts the SFS towards lower frequencies but with high abundance when *N* increases. Hence, a larger population size results in a smaller mean of minor allele frequencies and a wider variance of minor allele frequencies (Figure S1).

#### Genetic drift threshold

Alleles present at a very low frequency in the initial population are easily lost by genetic drift, independently of their selective advantage. Since we are interested in studying the population’s response to selection, we impose a genetic drift threshold to the frequency of the alleles, which ensures a lower probability that selected alleles are eliminated by drift at the beginning of the evolutionary process. Assuming a haploid population of size *N*, an *i*-copy minor allele has a frequency *i/N*. The probability of losing this minor allele in the following generation is *p*_lost_ = (1 − *i/N*)^*N*^. When *N* is large enough, this is *p ≈ e*^−*i*^, which is always true in our study since the smallest considered population size is 100.

Assuming we want to ensure that *p*_lost_ *<* 0.05, then *i* ≥ 3, i.e., we should consider minor alleles with at least three copies in the initial population. We define this as the genetic drift threshold in the population. With or without genetic drift, the mean of the minor allele frequency is correlated with the population size *N*, which spans 100 to 5000 in our simulations. The mean allele frequency is always below 0.2; it is largest for *N* = 100 and drops to below 0.1 as the population size increases (Figure S1A and S1B). Using the drift threshold slightly elevates the mean and variance of minor allele frequencies (Figure S1C and S1D), resulting in a more diverse initial population.

The drift threshold aims to reduce the initial effect of genetic drift in our simulations in the interest of efficient simulation. Without a threshold, a large number of alleles at very low frequencies become fixed immediately at the beginning of our simulation. Thus, we would have to study a more high-dimensional fitness landscape to observe the same dynamics of selection on the fitness landscape as we observe with the drift threshold. With the drift threshold, we focus on loci potentially contributing to fitness, resulting in a smaller fitness landscape to search and hence greatly improving the simulation efficiency.

#### Fraction of the fitness landscape covered by the initial population

When studying adaptation from SGV only, the initial state of the population plays a prominent role. Unless the fitness landscape is very small and the population is large, only a fraction of all genotypes in the genotype space will be represented in the initial population. We compute the expected initial occupation of the fitness landscape with *L* loci. The probability of an individual carrying a given genotype with *k* minor alleles is approximately 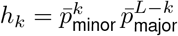. Thus, the probability that a specific genotype is not present in the population is (1 − *h*_*k*_)^*N*^. We can then sum the probability of a genotype being present in the population over all genotypes and divide this sum by the total number of possible genotypes. This corresponds to the expected fraction of the genotype space occupied by the initial population,

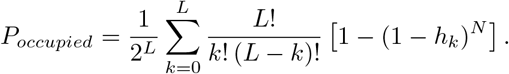

The factor after the sum accounts for the number of genotypes that contain *k* minor alleles. Given a genetic drift threshold, we can re-estimate *P*_*occupied*_ by setting the lower boundary as *i*_*t*_ = 3 (Figure 2).

**Figure 2.**
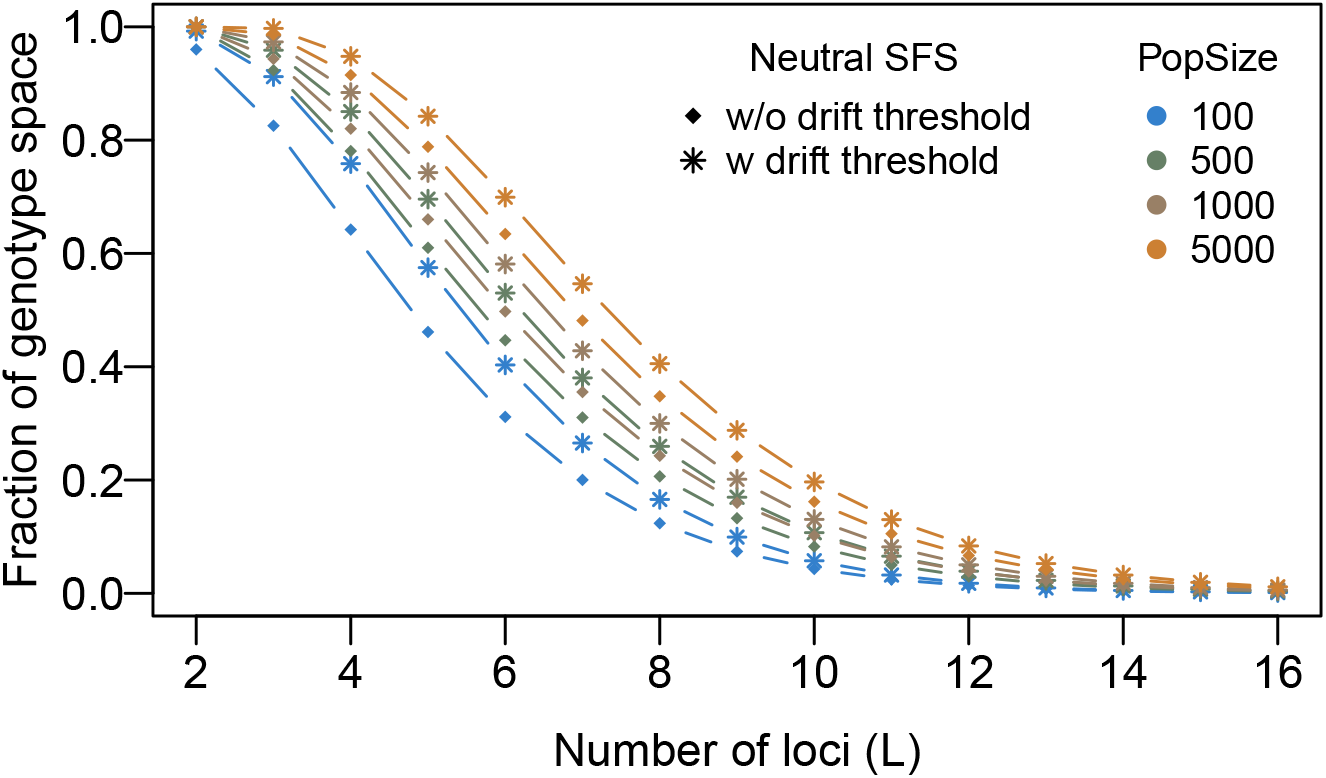
Approximate fraction of the genotype space initially covered by differently sized populations. Filled diamonds correspond to populations generated without (w/o) the drift threshold, stars correspond to populations generated with (w) the drift threshold. Four population sizes (PopSize) are shown. As the dimensionality of the fitness landscape increases, a smaller fraction of genotypes is covered by the initial population. The drift threshold increases the fraction of the genotype space covered because it increases the mean initial allele frequency. For the analytic expression used to plot the figure, see Methods.

#### Sampling a neutral population with the drift threshold

We sample the initial population with *N* individuals from a population in mutation-drift balance before changing the environment. Each genotype contains *L* loci. For each locus, we draw an integer *k* in the range *i* to *N − i*, distributed according to the probability 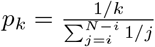, where *i* is the drift threshold. If the sampled integer is smaller than *N/*2, we assign allele 1 to that locus in *k* random individuals of the population. Otherwise, we assign allele 1 to *N* − *k* individuals at the focal locus. This strategy guarantees that we draw the population from a neutral SFS with a drift threshold *i* and that the minor alleles are always assigned the beneficial effect. We used a drift threshold of *i* = 3 throughout this study. Since we assign the minor alleles to random genotypes (individuals), assignments are independent of each other. Therefore, the initial population is at approximate linkage equilibrium.

### Selection along the adaptive path

In order to represent empirical cases in which the identity and order of fixed alleles may be known, but a characterization of the combinatorially complete fitness landscape is rarely possible, we extract the direct selection coefficients and pairwise epistasis along the observed adaptive trajectory in the fitness landscape (Figure S2). It is important to note that there is no “adaptive walk” in the true sense since the population is polymorphic until the end of the simulation; we thus use the term “adaptive path” throughout the paper to denote the successive fixation of alleles. During evolution on a fitness landscape with *L* initially segregating loci, *L* fixation events happen on the adaptive path, and we refer to each fixation event as a “fixation step”. When two fixation events happen simultaneously, these are accounted for as separate fixation steps in our statistics, with their order randomized.

As illustrated in Figure S2, we define either the final genotype or the initial major genotype as a reference genotype that represents the genetic background. We obtain the one-locus effect as the effect of a single substitution between the fixed and the lost alleles at each fixation step. Here, the remaining *L* − 1 loci are assumed to be fixed for the reference alleles. We infer pairwise epistatic effects as the two-locus epistasis that is measured when considering two consecutive fixation steps, such that the (sub-)fitness landscape spans four genotypes. We extract the pairwise epistatic coefficient of the genotype with two fixed alleles when the remaining *L* − 2 loci are assumed to be fixed for the reference alleles. For example, the pairwise epistatic coefficient is *f*_001_ + *f*_100_ − *f*_101_ − *f*_000_ when the first (allele 0) and the third (allele 1) locus are fixed consecutively (Figure S2) and the reference allele at locus 2 is 0.

### Data Availability

Data sharing is not applicable to this article as no empirical datasets were generated or analysed during the current study. All annotated simulation code to reproduce the study will be archived on Zenodo upon publication of the paper, and the doi will be provided here.

## Results

### Initial occupation of the fitness landscape

Diallelic fitness landscapes are hypercubes of size 2^number of loci^. Thus, the number of genotypes in these landscapes increases rapidly as the number of loci increases.

Consequently, even if every locus is originally polymorphic, only a fraction of possible genotypes can be present in a population unless the fitness landscape is very small. Without *de novo* mutation, recombination is necessary to create any additional genotypes.

Based on the mean minor allele frequency, we computed the number of genotypes in the initial population given an *L*-locus genotype (Figure 2; see Methods). Here, the smallest landscape contains 2^2^ = 4 genotypes, whereas the largest reaches 2^16^ = 65536 genotypes. Hence, already for a moderate number of loci, the size of the genotype space grows much larger than the population size used in our simulations (*N* ∈ [100, 500, 5000], Figure 2). An initial population with SGV only covers the whole genotype space when the fitness landscape is very small. When the fitness landscape has many loci, a population can only cover a small fraction of genotypes in the genotype space. In these larger landscapes, nearly every individual is unique, and each genotype is mainly composed of major alleles.

### Time to fixation is affected subtly by the recombination probability

In our simulations, adaptation occurs until the standing genetic variation in the population is exhausted and the population becomes monomorphic. The required time to fixation of a single genotype depends on a range of factors including the ruggedness of the fitness landscape, the population size, and the recombination probability.

The ruggedness of the fitness landscape greatly influences the fixation time. Figures 3A and 3B show that populations achieve fixation one order of magnitude faster in the case of highly rugged landscapes relative to fitness landscapes with low or intermediate ruggedness. Three factors contribute to this pattern. Firstly, higher ruggedness entails stronger selection between genotypes since the fitness differences between neighboring genotypes are larger, which is partly due to our parameterization of the RMF model. Secondly, highly rugged fitness landscapes have many local peaks at which the population can get trapped. Thirdly, the global peak is located (almost) randomly with respect to the major genotype in a highly rugged fitness landscape, whereas it is by definition located (approximately) “opposite” of the major genotype in a smooth fitness landscape. The global peak is therefore, on average, more strongly represented in the initial population in a highly rugged than in a smooth fitness landscape. The third factor is a consequence of our definition of the initial population distribution, which does not affect our results qualitatively (not shown). These features shorten the time until the population becomes monomorphic in highly rugged fitness landscapes. Because the location of the peaks is random, higher ruggedness also leads to a wider variation in fixation times. We are unable to tear apart how much of the absolute difference in different statistics (such as the fixation time) between ruggednesses is a result of the definition of the RMF model *versus* how much is a consequence of epistasis *per se*. Therefore, we mainly focus on the effect of different recombination probabilities on the statistics throughout the rest of the paper.

**Figure 3.**
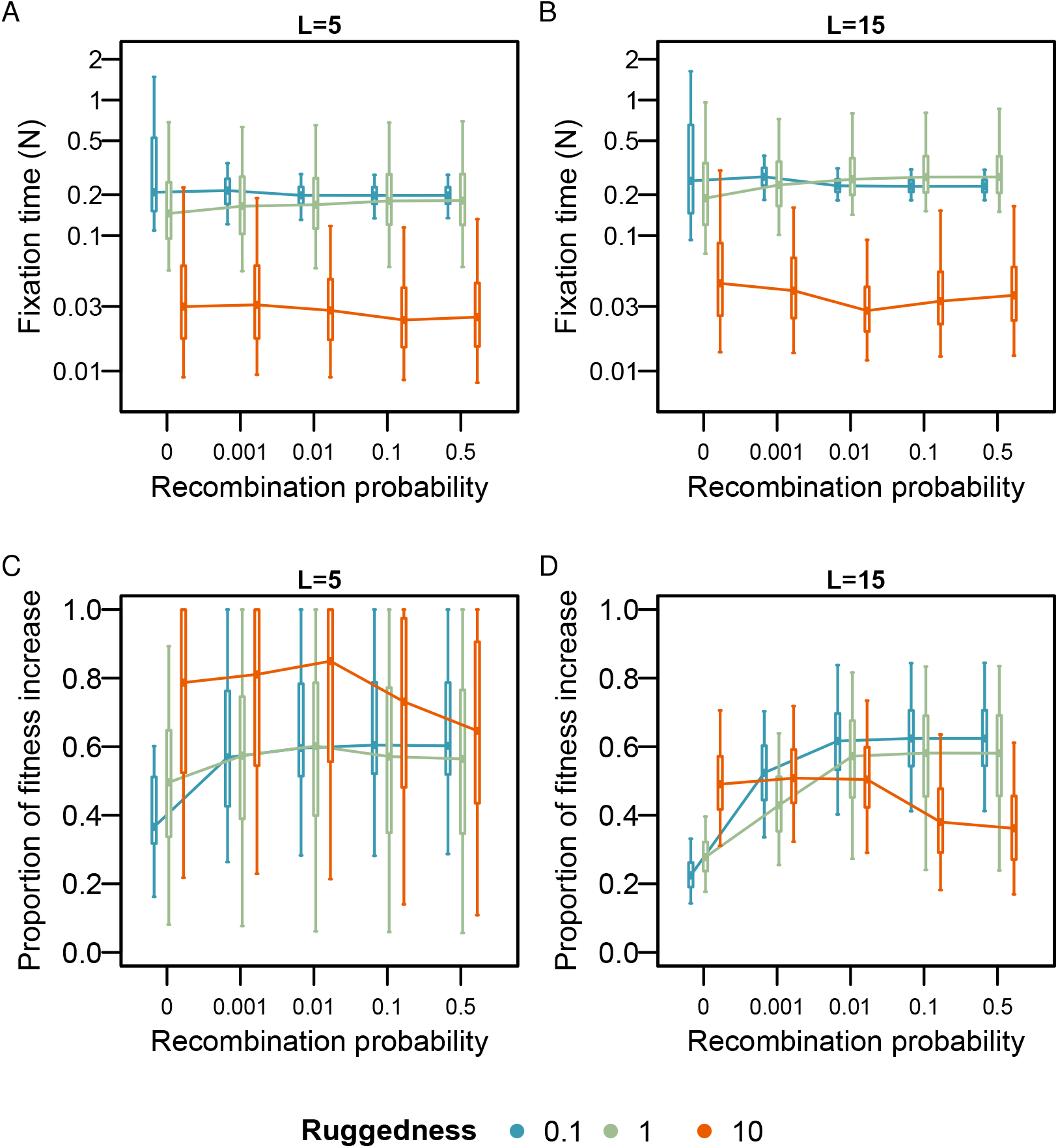
Fixation time and relative fitness change. A & B. Times until the population becomes monomorphic, scaled by the population size. C & D. Fitness change between the initial mean fitness and the fitness of the final genotype, normalized to the distance of the mean initial population fitness to the global peak. For both small (A; *L* = 5) and large (B; *L* = 15) fitness landscapes, fixation occurs rapidly in rugged fitness landscapes. Although the fixation time is similar across recombination probabilities, the fitness gain through adaptation decreases with the recombination probability for highly rugged fitness landscapes (C & D), whereas it increases with the recombination probability for smooth and intermediately rugged, large fitness landscapes (D). For each parameter set, we simulated 100 fitness landscapes and allowed 100 populations with different initial genotype distributions to adapt on each fitness landscape. Each population contains 5000 individuals (*N* = 5000). We study fitness landscapes of varying ruggedness (0.1, 1, 10) by studying three epistasis parameters, 0.001, 0.01, and 0.1, while maintaining a constant additive effect 0.01. Boxes represent the interquartile range, and whiskers extend outwards to cover 95% of the data. Statistical p-values are provided in Table S1. See also Methods.

On large and smooth fitness landscapes, fixation times decrease with increasing recombination probability, whereas the fixation time increases with recombination probability in large and intermediately rugged fitness landscapes. For large and highly rugged landscapes, we observed a minimum of the fixation time and its variance at intermediate recombination probabilities. These effects are systematic but weak compared to the variation among replicates (see p-values in Table S1). The variance in the fixation time is always largest when there is no recombination, a trend that is especially visible for smooth fitness landscapes. The population size also plays a crucial role in determining the fixation time, with larger populations reaching fixation relatively faster than smaller populations, measured in units of the population size (Figure S3). In general, the effects of population size (and landscape ruggedness, but see above) on the fixation time are much larger than the effects of landscape size and recombination probability.

### Fitness gain decreases with increasing recombination at high ruggedness

Recombination breaks the linkage among positively epistatic alleles (*e*.*g*., Neher and Shraiman 2009). Thus, high recombination should reduce the potential gain of mean population fitness in epistatic fitness landscapes. On the other hand, recombination brings together beneficial alleles in additive landscapes, thereby reducing Hill-Robertson interference (HRI; Hill and Robertson 1966). We calculated the fitness change from the initial mean population fitness to the final genotype, normalized by the distance of the mean fitness of the initial population to the global fitness peak. (I.e., if the fitness change is one, the population evolved to the global fitness peak.) As expected, in smooth fitness landscapes, the fitness gain increases with the recombination probability (*λ* = 0.1 or 1, Figure 3 and S4). Here, increasingly many beneficial mutations (i.e, those with an additive beneficial effect) are fixed because recombination reduces HRI. We observe the reverse pattern in highly rugged fitness landscapes. Here, the gain in population fitness decreases with increasing recombination probability (Figure 3, S4C and S4F).

### Initial frequencies determine which alleles become fixed on rugged fitness landscapes

We assigned the positive additive effect to minor alleles; therefore, if direct selection is effective, minor alleles should predominantly become fixed, resulting in increased divergence from the initial population. Correspondingly, divergence increases with recombination probabilities for fitness landscapes with low and intermediate ruggedness (*λ* = 0.1 or 1, Figure 4A, 4B and S5). Here, high recombination probabilities favor the fixation of genotypes with more beneficial (minor) alleles (Figure 4C-E, S6 and S7).

**Figure 4.**
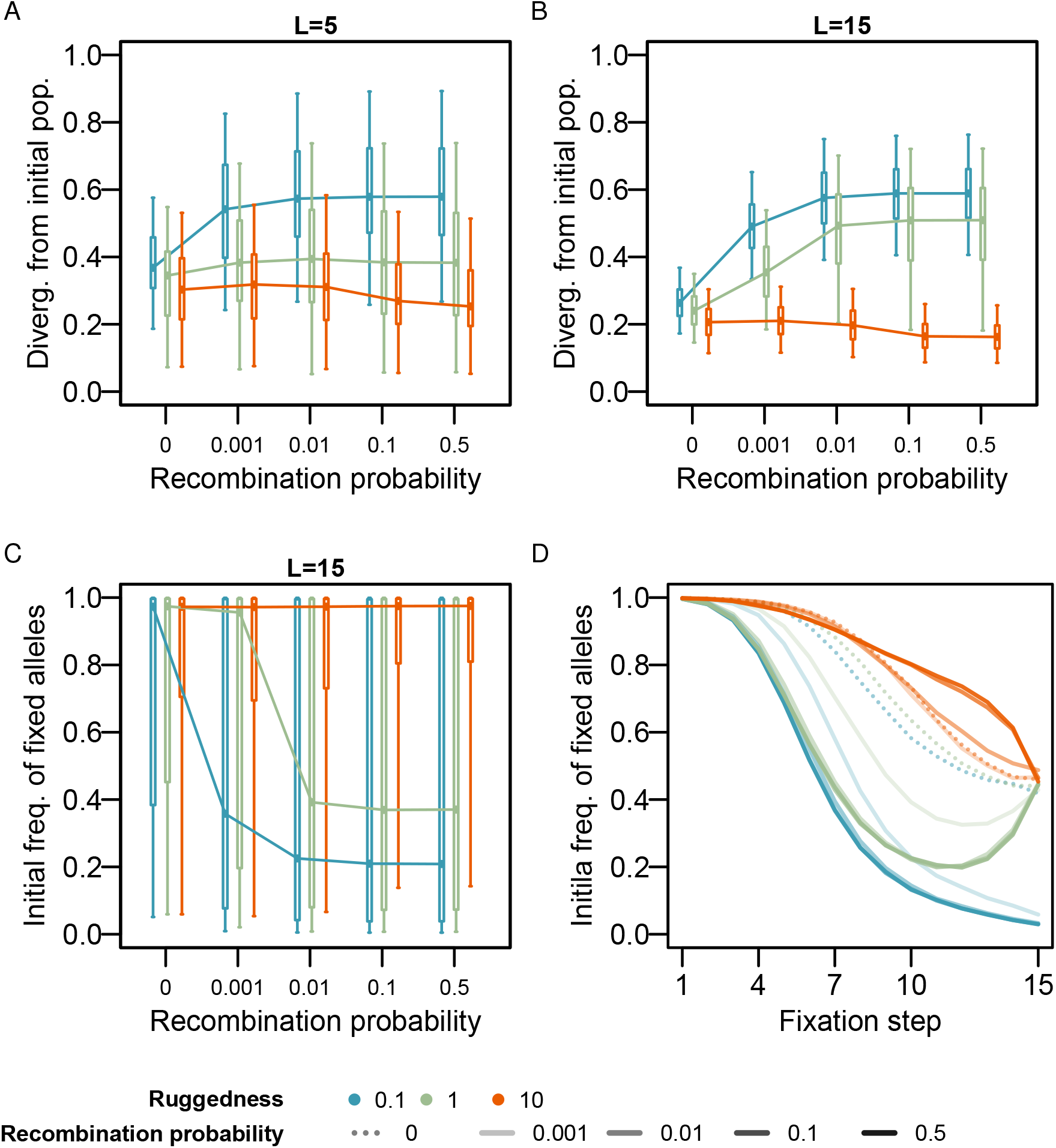
Genetic change between the initial population and the final genotype. To measure the divergence between the initial and the final populations, we calculated the average normalized Hamming distance between the final genotype and the initial population. A. Divergence for small fitness landscapes (*L* = 5). B. Divergence for large fitness landscapes (*L* = 15). The final evolved genotype is closer to the initial population when both ruggedness and recombination probabilities are high. For low and intermediate ruggedness, the divergence from the initial population increases with the recombination probability. C & D show the initial frequencies of the fixed alleles (*L* = 15). C. The mean initial frequency of fixed alleles at four recombination probabilities. D. The mean initial frequency of fixed alleles along the adaptive path. Each fixation step corresponds to a fixation event on the adaptive path. More high-frequency alleles are fixed by the stronger interplay of recombination and epistasis in rugged fitness landscapes. Boxes represent the interquartile range, and whiskers extend outwards to cover 95% of the data. Statistical p-values are provided in Table S1. All other simulation details are as in Methods and Figure 3.

When the fitness landscape is highly rugged, divergence from the initial population decreases with recombination (*λ* = 10, Figure 4A and 4B). At high ruggedness, we observe that the initial frequency of fixed alleles is exceedingly high and increases with recombination probability (*λ* = 10, Figure 4C, S6 and S7). We suspect that a high recombination probability favors the fixation of combinations of positively epistatic high-frequency alleles. Moreover, at high ruggedness, high-fitness genotypes are more evenly distributed across the fitness landscape than in smoother landscapes since fitness is mainly determined by epistatic combinations of alleles instead of the direct effects of the alleles. Thus, the chance that the initial population contains high-fitness genotypes, including the global peak, at appreciable frequencies is high in highly rugged fitness landscapes. Notably, the global peak is almost never the final genotype in large fitness landscapes of any ruggedness (Figure S8).

Across all epistasis parameters, mostly major alleles become fixed during the first few steps of the adaptive path (Figure 4D and S6). This can likely be attributed to genetic drift and competition among many selected genotypes. During these initial fixation steps, individually beneficial (minor) alleles are lost even in the presence of recombination. At low ruggedness, the mean initial allele frequencies of fixed alleles over the fixation trajectory for different recombination probabilities separate very early, indicating that recombination begins to prevent beneficial alleles from being lost (step 2, Figure 4D and S6). Therefore, at high recombination probabilities, more minor (beneficial) alleles become fixed over time (Figure 4D). At high ruggedness, the mean initial allele frequencies of fixed alleles over the fixation trajectory remain indistinguishable until fixation step 7 (Figure 4D). In highly rugged fitness landscapes, many high-fitness genotypes are distributed randomly across the fitness landscape, and hence there is strong competition among many selected genotypes. Because several of these genotypes may exist initially at appreciable frequencies, selection can act on them early during the adaptive path (Figure S9). Hence, high-frequency alleles involved in positively epistatic genotypes become fixed rapidly.

Populations that evolve on intermediately rugged fitness landscapes (*λ* = 1) follow a U-shaped fixation pattern. During the first steps of the adaptive path, the initial frequency of fixed alleles decreases, following the pattern of smooth fitness landscapes (Figure 4D and S6). Alleles with initially higher frequency become fixed towards the end of the adaptation (Figure 4D and S1). At the final fixation step, the mean initial frequency of the fixed allele for intermediate ruggedness converges to the value for high ruggedness, where fixation is random. At this step, selection in intermediately rugged fitness landscapes mainly comes from epistasis (step 15, Figure 5A middle panel).

**Figure 5.**
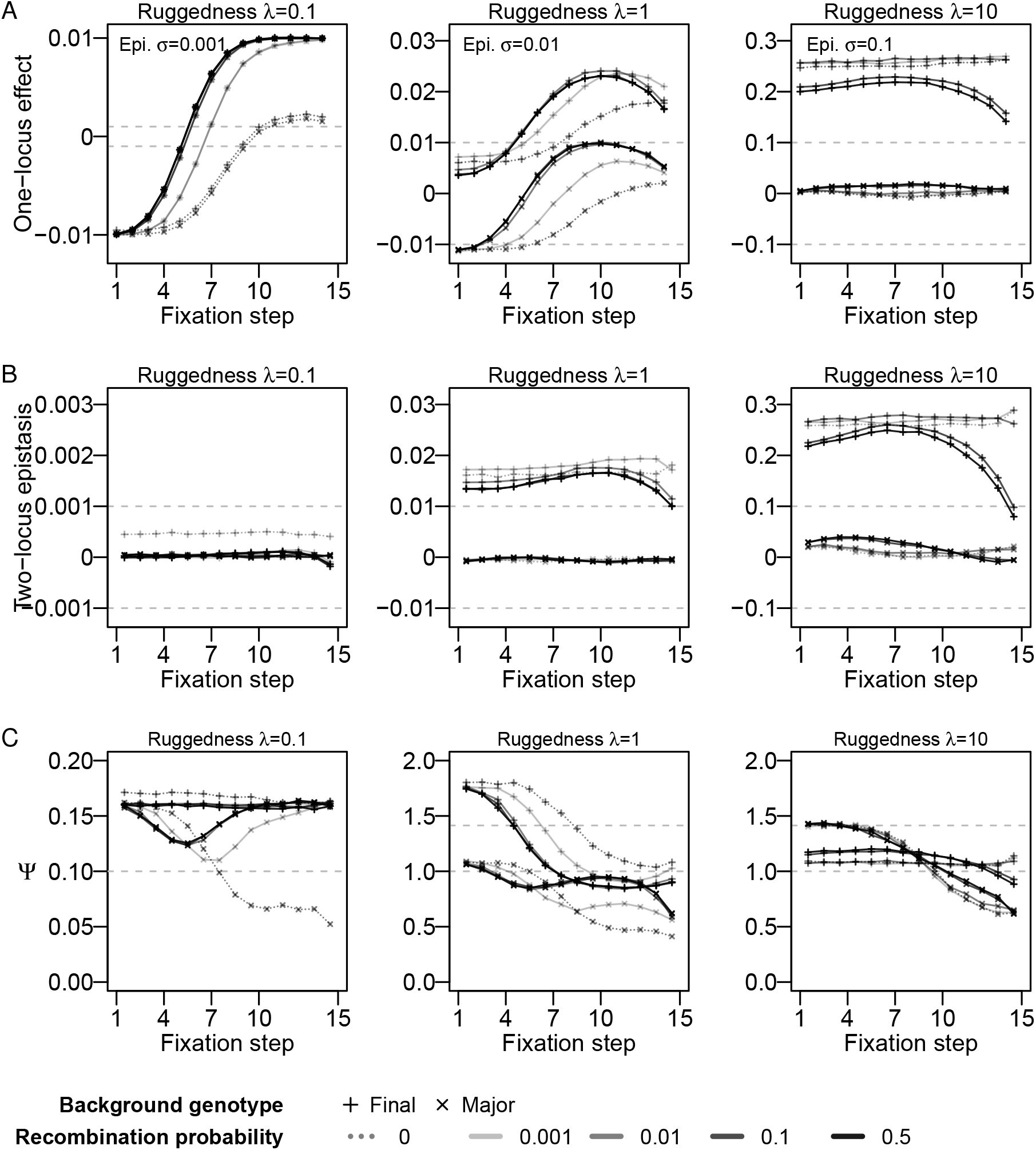
Selection coefficients extracted from the adaptive path. We extracted the one-locus effect (A) and the pairwise epistasis (B) based on two monomorphic genetic backgrounds: the initial major genotype (crosses) and the final fixed genotype (pluses). Here, the one-locus effect is the selection coefficient of the fixed allele compared to the lost allele at each fixation step, where all other loci are fixed for the focal genetic background. Pairwise epistasis is the epistasis observed between two consecutive fixed alleles on the focal background. Adaptation of polymorphic populations occurs along a smoother-than-average path in the fitness landscape: one-locus effects are larger, and epistasis is lower than expected from the underlying fitness landscape. The ruggedness parameter is indicated above each panel, and the epistatic standard deviation (*σ*) is highlighted by dashed lines in each panel for scale. Note the different y-axis ranges between panels. (C). The ratio of mean pairwise epistasis and absolute one-locus effect Ψ describes the observed ruggedness along the adaptive path. Dashed lines represent ruggedness (0.1, left panel), Ψ values of 1 and 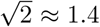 (middle and right panels). Simulation details are as in Methods and Figure 3.

### Weak epistasis and strong direct selection on the adaptive path

When adaptation from SGV is observed in nature, rebuilding the complete underlying fitness landscape experimentally is generally not feasible. Even extracting two-locus fitness landscapes is a large experimental endeavor, but it is more realistic and has been achieved occasionally. Here, we use our simulation data to extract one-locus selection effects and pairwise epistasis along the observed adaptive paths, thereby showing which estimates would be obtained in experimental tests of pairwise subsets of a fitness landscape sampled along an adaptive path. We define as reference genetic background either the final fixed genotype or the major genotype, i.e., the genotype composed of the major alleles at each locus. We then measure the one-locus effects of the fixed alleles and the pairwise epistasis between two consecutively fixed alleles (see Methods, Figure S2).

In general, we cannot extrapolate from the extracted direct selection and pairwise epistasis along the adaptive path to the shape of the underlying fitness landscape. When the landscape is smooth, the extracted one-locus selection coefficient increases from -0.01 to 0.01 with the sequential fixation of alleles (Figure 5A, left panel), independent of which genotype is chosen as the reference. This pattern reflects that initially, genetic drift fixes the major, deleterious allele. Towards the end of the adaptive path, the direct selection coefficient reflects its corresponding model parameter, except when the recombination probability is low or zero. For smooth fitness landscapes, we observe consistently lower pairwise epistasis along the adaptive path than in the underlying landscape (Figure 5B, left panel).

When the fitness landscape is intermediately or highly rugged, the choice of reference genotype greatly affects the extracted selection and epistasis along the adaptive path (Figure 5, S10 and S11A). Taking the final genotype as the reference, the extracted one-locus effects and pairwise epistasis along the adaptive path are larger than expected from the underlying fitness landscape. In contrast, when the major genotype is chosen as the reference, the extracted one-locus effects and pairwise epistasis along the adaptive path are smaller than expected from the underlying fitness landscape. In general, the extracted one-locus effects along the adaptive path are of the order of the epistasis (standard deviation of the epistatic contribution *σ*). The amount of pairwise epistasis extracted along the adaptive path on the basis of the final genotype is of the order of the epistasis *σ*, whereas almost no pairwise epistasis is observed along the adaptive path when the major genotype is chosen as a reference.

Near-zero one-locus effects and two-locus epistasis values in the major genotype background could originate from either very small effects or cancellations due to averaging. To distinguish between these two scenarios, we computed the means of the absolute values of the effects on both backgrounds (Figure S10 and S11). Both effects are of the order of *σ* when the final genotype is taken as a reference, indicating that during adaptation positive direct selection and epistasis are accumulated in the final genotype background. In contrast, taking the major genotype as a reference, the mean of the absolute values is on the order of *σ*, which shows that here, the near-zero mean values reported above arise from cancellations of positive and negative effects, representing the random initial condition in our simulations.

The recombination probability also influences the observed selection and pairwise epistasis along the adaptive path. Low recombination probabilities lower the extracted one-locus effects for smooth and intermediately rugged fitness landscapes. In contrast, low recombination probabilities increase the observed one-locus effects along the adaptive path for very rugged 2 *≈* 1.4 (middle and right panels). Simulation details fitness landscapes, when the final genotype is chosen as a reference. Moreover, the observed values vary more between the beginning and the end of the adaptive path at high recombination probabilities. For example, the extracted pairwise epistasis along the adaptive path, obtained with the final genotype as a reference, approximately reflects the epistasis in the underlying fitness landscape for the last pair of fixed mutations for intermediately and highly rugged fitness landscapes when the recombination probability is high (*r* = 0.1 or *r* = 0.5; Figure 5, right panel). In contrast, the extracted pairwise epistasis along the adaptive path takes rather different values and remains almost constant for lower recombination probabilities.

We observed similar patterns for small population sizes (*N* = 100) and small fitness landscapes (*L* = 5, Figure S10 and S11). Notably, strong genetic drift in small populations leads to generally lower observed one-locus selection coefficients and pairwise epistasis along the adaptive path, except in highly rugged fitness landscapes. Moreover, for fitness landscapes of intermediate and large ruggedness, the variation in observed one-locus effects and pairwise epistasis between the beginning and the end of the adaptive path, and between reference genotypes, is lower when the fitness landscape is small (*L* = 5, Figure S10 and S11).

### Adaptation proceeds along a smoother-than-average trajectory on highly-rugged fitness landscapes

We reported above that epistasis in the fitness landscape is observed as direct selection along the adaptive path when taking the final genotype as a reference. To quantify whether this observation is specific to the adaptive path, we computed the ratio of the means of the absolute value of pairwise epistasis and the absolute one-locus effect

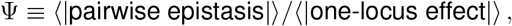

where | *·* | denotes the absolute value and ⟨·⟩ the mean value, as a proxy of the ruggedness of the experienced fitness landscape, similar to the definition of the ruggedness of fitness landscapes (see Methods; Aita et al. 2000). A high value of Ψ reflects that the population experiences a rugged fitness landscape, whereas a value close to zero implies that the population is moving through a smooth region of the fitness landscape.

To develop an expectation of Ψ, we computed the expected value of Ψ for a random walk in the House-of-Cards model (the high-ruggedness limit of the RMF model, where genotype fitness is random). Pairwise epistasis depends on four genotypes (*e*.*g*., *f*_11_ − *f*_01_ − *f*_10_ + *f*_00_, Figure S2) and the one-locus effect on two genotypes (*e*.*g*., *f*_01_ − *f*_00_, Figure S2).

Consequently, we expect their values to be distributed normally with standard variation 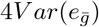 and 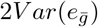, respectively. Knowing these distributions and using the expected absolute value of pairwise epistasis and expected absolute value of one-locus effects, the expected value of Ψ in the House-of-Cards limit is

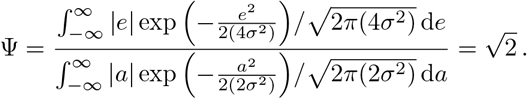

The expected value of 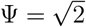 for the House-of-Cards model agrees well with the Ψ observed for our highly rugged RMF landscapes (*λ* = 10; Ψ = 1.410 *±* 0.001, obtained from 10^6^ two-locus RMF fitness landscapes).

The opposite limiting condition is no epistasis (Ψ = 0). Between the two limiting conditions, we simulated three levels of ruggedness (*λ* = 0.1, 1, 10). On smooth (*λ* = 0.1) and intermediately-rugged fitness landscapes (*λ* = 1), the Ψ expectations are 0.1594 *±* 0.0001 and 1.140 *±* 0.001 (averaged over 10^6^ two-locus RMF fitness landscapes for each). Along adaptive paths in our studied fitness landscapes, when the major genotype is taken as a reference, Ψ is usually lower than expected, but Ψ is approximately equal to or larger than expected when the final genotype is taken as a reference (*λ* = 0.1&1, Figure 5C and S12). This indicates that adaptation ignores epistasis or favors regions with some epistasis, when epistasis contributes weakly to the fitness landscapes. At high ruggedness, we observe that the population moves along an adaptive path towards lower ruggedness. Whereas initially, when the reference is the major genotype, the ratio Ψ is close to the expectation of 1.4, it decreases to values lower than one during the adaptive path (Figure 5C and S12). When the reference is the final genotype, the ratio Ψ is generally lower than when the reference is the major genotype, and it also decreases along the adaptive path. This indicates that populations take a path from a random location in the fitness landscape to regions of lower ruggedness.

## Discussion

Probabilistic fitness landscape models are a playground to study the effect of epistasis on adaptation. Here, we study rapid adaptation from standing genetic variation of recombining haploid populations on fitness landscapes of varying ruggedness, using realistic initial population compositions. The role of epistasis in adaptation and its interaction with recombination is a challenging topic, and our study illustrates the complex effects of recombination on adaptation in the presence of epistasis. We further show that adaptation proceeds along smoother-than-average paths and that rugged fitness landscapes could leave a signal of strong direct selection in empirical data.

Both theoretical and empirical studies have highlighted the crucial role of standing genetic variation for adaptation to environmental challenges (e.g., Barrett and Schluter 2008; Orr and Unckless 2014; Lai et al. 2019; Marques et al. 2019; Bomblies and Peichel 2022). Once adaptation has occurred, it is a challenge to infer how much epistasis and direct selection, respectively, contributed to the adaptive path. This is an important question because the result indicates whether adaptation will be repeatable on different genetic backgrounds in other populations exposed to similar environmental challenges. If direct selection is primarily responsible for adaptation, the same alleles will contribute to adaptation repeatedly. If epistasis plays a dominant role, adaptation is contingent on the genetic background and different genotypes will be spreading in different populations. Knowing which of these scenarios is at play is crucial for guided interventions in conservation biology (so-called genetic rescue), for example, because epistasis could render the introduction of genetic material ineffective or even detrimental.

In our study, polygenic adaptation mostly occurs within 1000 generations. This time scale of rapid adaptation is similar to what has been described in polygenic human adaptation, *e*.*g*., regarding the increased height of northern Europeans (Turchin et al. 2012). It was estimated that there are as many as *>*1000 “tall” alleles that have been segregating under weak selection (selection coefficients ∼ 10^−3^ − 10^−5^ per allele, Turchin et al. 2012). Weak selection has increased their frequency in the past 2000 years (*<* 100 generations) (Field et al. 2016).

Another example of rapid polygenic adaptation comes from plant traits of sedge (*Schoenoplectus americanus*) that rapidly responded to sea level rise within 50 years, and the corresponding genotypes are heritable (Vahsen et al. 2023). In our study, direct selection with selection coefficient *a* = 0.01 per allele could fix a set of standing genetic variants within 1000 generations regardless of population size (Figure 3 and S3). When epistasis brings associations among loci, polygenic adaptation was even faster, fixing all variants in less than 200 generations (Figure 3 and S3).

On additive fitness landscapes, recombination is necessary to bring together beneficial alleles that segregate on different genetic backgrounds. When a locus is under strong direct selection, fixation probability and time are correlated negatively (Kimura and Ohta 1969). For multiple loci, high recombination probabilities increase the fixation probability and accelerate fixation by combining beneficial alleles in a single genotype (Figure 3A&B, Roze and Barton 2006). At first sight, surprisingly, we find almost no effect of the recombination probability on the time until fixation of a monomorphic genotype (Figure 3A&B). However, this result is explained when taking into consideration the divergence from the initial genotype and the fitness gain during adaption at different recombination probabilities (Figure 3C&D and 4A&B). When there is no recombination, no new genotypes can be created, and fixation occurs on average at lower fitness and with less average divergence from the initial genotype composition. Therefore, although the adaptive path is “shorter”, the fixation time is similar across recombination probabilities. Thus, in agreement with the abovementioned classical results, adaptation on smooth fitness landscapes occurs more slowly and less effectively at low recombination probabilities.

Epistasis is a type of second-order selection that is affected by recombination in a more complex fashion than direct selection. On the one hand, epistasis is only expressed when different allelic combinations segregate simultaneously (i.e., recombination or high mutation rates are required to create the different haplotypes); on the other hand, recombination breaks apart positively epistatic combinations of alleles, thereby reducing the selection effect of epistasis. The positive frequency-dependent nature of epistasis drives alleles at epistatically interacting loci towards fixation (Ayala and Campbell 1974). In our study, we initialize populations at linkage equilibrium. Due to the shape of the SFS, many alleles are initially at very high or low frequencies and many high-fitness genotypes are present initially in highly rugged fitness landscapes. This explains our observation that, in highly rugged fitness landscapes, a high recombination probability (*r* = 0.1 or *r* = 0.5) is detrimental to fitness gain during adaptation.

Consistent with previous reports of an “optimal” recombination rate for crossing a two-locus fitness valley (Weissman et al. 2010), we observed small-effect but systematic evidence for a minimum time to fixation while maintaining an optimal fitness gain for an intermediate recombination probability in large and highly rugged fitness landscapes (*λ* = 10 and *L* = 15; see Figure 3B&D). Similarly, for small and highly rugged fitness landscapes, the same intermediate recombination probability seems to result in a maximum fitness gain while maintaining an almost constant fixation time across all recombination probabilities (*λ* = 10 and *L* = 5; see Figure 3A&C). This pattern is also visible for small, intermediately rugged fitness landscapes, in which direct selection and epistasis are of the same order of magnitude (*λ* = 1 and *L* = 5; Figure S4). However, in large, intermediately rugged fitness landscapes, increasing recombination is always beneficial for fitness gains (*λ* = 1 and *L* = 15; Figure S4). We suspect that the added value of recombination, by bringing multiple alleles of additive beneficial effect into the same genotype, overrides the intermediate optimum for large fitness landscapes. Here, a smaller proportion of genotypes in the fitness landscape exist in the initial population (and at lower frequencies, which reduces the effectiveness of epistasis).

Notably, the divergence from the initial population increase with recombination probability on smooth fitness landscapes but decreases on highly rugged landscapes. That is because the number of fitness peaks is large in highly rugged fitness landscapes; therefore, (high-fitness) peaks are distributed across the fitness landscape (Bank 2022). This increases the probability of high-fitness genotypes being present in the initial population. Therefore, in highly rugged fitness landscapes, a significant fitness increase can often be achieved without recombination (Figure 3C&D). On the other hand, populations that evolve in large and highly rugged fitness landscapes are in danger of getting stranded at suboptimal genotypes, especially if there is strong recombination, which prevents the establishment of newly recombined high-fitness epistatic genotypes. Previous work studying the evolution of recombining populations in fitness landscapes by means of *de novo* mutations has described a similar pattern: when recombination is strong, populations can get stalled at a suboptimal peak for much longer than with intermediate recombination (de Visser et al. 2009; Nowak et al. 2014).

On highly rugged fitness landscapes, the larger the product of genotype frequency and its pairwise epistatic interaction *e*_*g*_, the more likely a genotype becomes fixed. Imagine an extreme example in which alleles *A* and *B* interact positively (*e*_*AB*_ *>* 0) and all other genotypes (*Ab, aB, ab*) are neutral. Genotype frequency is denoted as *x*. If *x*_*AB*_ * *e*_*AB*_ *> x*_*ab*_ * *x*_*AB*_ * *r*, the positive epistasis between allele *A* and *B* can overcome the breakdown effect by recombination *r*. Subsequently, *AB* is driven towards fixation and its spread is accelerated when *AB* becomes common in the population. This type of genotype selection is known as “clonal condensation” (Neher et al. 2013). In Neher et al. 2013‘s work, populations were initialized as a maximally diverse sample at linkage equilibrium, such that each allele was at an equal frequency (around 0.5) and each genotype was equally at a low frequency (predominantly 1 copy). The populations subsequently evolved via selection and recombination. Once some genotypes reached high frequencies, a clonal structure emerged at high recombination rates on highly-rugged fitness landscapes (Neher et al. 2013). This is counterintuitive but similar to the positive frequency-dependent selection discussed above. “Clonal condensation” should be more pronounced in our study than in Neher et al. 2013 since our populations are initiated from a neutral SFS, where allele frequencies are biased to extreme values, such that genotype frequencies vary more than that in a maximally diverse population.

We showed here that inferring the one locus effects of fixed alleles and the pairwise epistasis between successively fixed alleles is not sufficient to quantify the shape of the underlying fitness landscapes. In line with our findings but using a rather different approach, Du Plessis et al. 2016 showed that regression models are often poor descriptors of the local fitness landscape when samples were chosen among adapted genotypes. Throughout the parameter range of our study, both the extracted direct selection and pairwise epistasis along the adaptive path do not reflect the parameters of the underlying fitness landscape. Specifically, on rugged fitness landscapes and when the final genotype is taken as the reference genetic background, we consistently observed one-locus effects on the order of magnitude of the epistasis parameter of the underlying fitness landscape, even if the true underlying additive effects were an order of magnitude smaller. Both the one-locus effect and pairwise epistasis along the adaptive path are also greatly dependent on the choice of reference genetic background. When the major genotype (i.e., the genotype with the initially most common alleles at each locus) is used as a reference, both effects vanish. From an empirical point of view, an important take-home message from this analysis is that inferring strong direct selection of recently fixed individual alleles could also be a signature of epistasis. Future work could take a genomic approach and test whether genomic signatures of epistasis are distinguishable from those of direct selection.

We observed that the local ruggedness decreases along the adaptive path on highly-rugged fitness landscapes. During the first fixation steps, we observe the highest ruggedness. Subsequently, selection and recombination weaken genotype competition by driving allele frequency from intermediate to extreme frequencies until fixation. Since genotype competition is strongest when many genotypes are evenly distributed, local ruggedness decreases when fewer genotypes segregate in the population because epistasis mimics direct selection. This observation differs from results obtained under the common assumption of strong selection and weak mutation, where new mutations arise and fix successively as an adaptive walk. These studies indicated that negative interactions tend to emerge early and positive epistasis later (Draghi and Plotkin 2013; Greene and Crona 2014; Blanquart et al. 2014). At first sight compatible with our findings, Blanquart et al. 2014 showed that subsets of a larger fitness landscape that are comprised of alleles that were fixed along an adaptive walk in Fisher’s Geometric model are, on average, smoother than subsets of a fitness landscape comprised of random alleles. However, in their model, the population adapts from smooth towards rugged regions on the genotypic fitness landscape. Here, as the population approaches the single phenotypic optimum, the same mutations that would have been beneficial far away from the optimum become deleterious (and, therefore, negatively epistatic) because they overshoot the optimum.

In this study, we modeled a haploid recombining population with standing genetic variation. We chose this simplification because dominance in diploid populations results in additional complexities, and existing fitness landscape models do not provide a definition of the dominance of additive effects and or the dominance of epistasis (Turelli and Orr 2000). Future work should address this problem by building sensible diploid models of fitness landscapes that include dominance in a parameter-light way. Moreover, the RMF model is neat for its simplicity but results in regular fitness landscapes. Although some experimental fitness landscapes were consistent with an underlying RMF landscape (Szendro et al. 2013; Bank et al. 2016), there is increasing evidence that real fitness landscapes are not as regular as the RMF model suggests (e.g., Pokusaeva et al. 2019). There are efforts to build fitness landscape models which account for irregularities of interactions (e.g., Reddy and Desai 2021) but these are not as easily studied and implemented as the RMF model, which requires only a single parameter to vary epistasis.

## Acknowledgements

We thank David McLeod, Stephan Peischl and the members from THEE division for suggestions that improved this work. We thank Wolfgang Stephan, Joachim Krug, and Georgii Bazykin for their constructive reviews and thoughtful comments during the review process. This work was supported by funding from ERC Starting Grant 804569 (FIT2GO), SNSF Project Grant 315230/204838/1 (MiCo4Sys), and HFSP Young Investigator Grant RGY0081/2020 to CB.

## Supplementary Information

**Figure S1.**
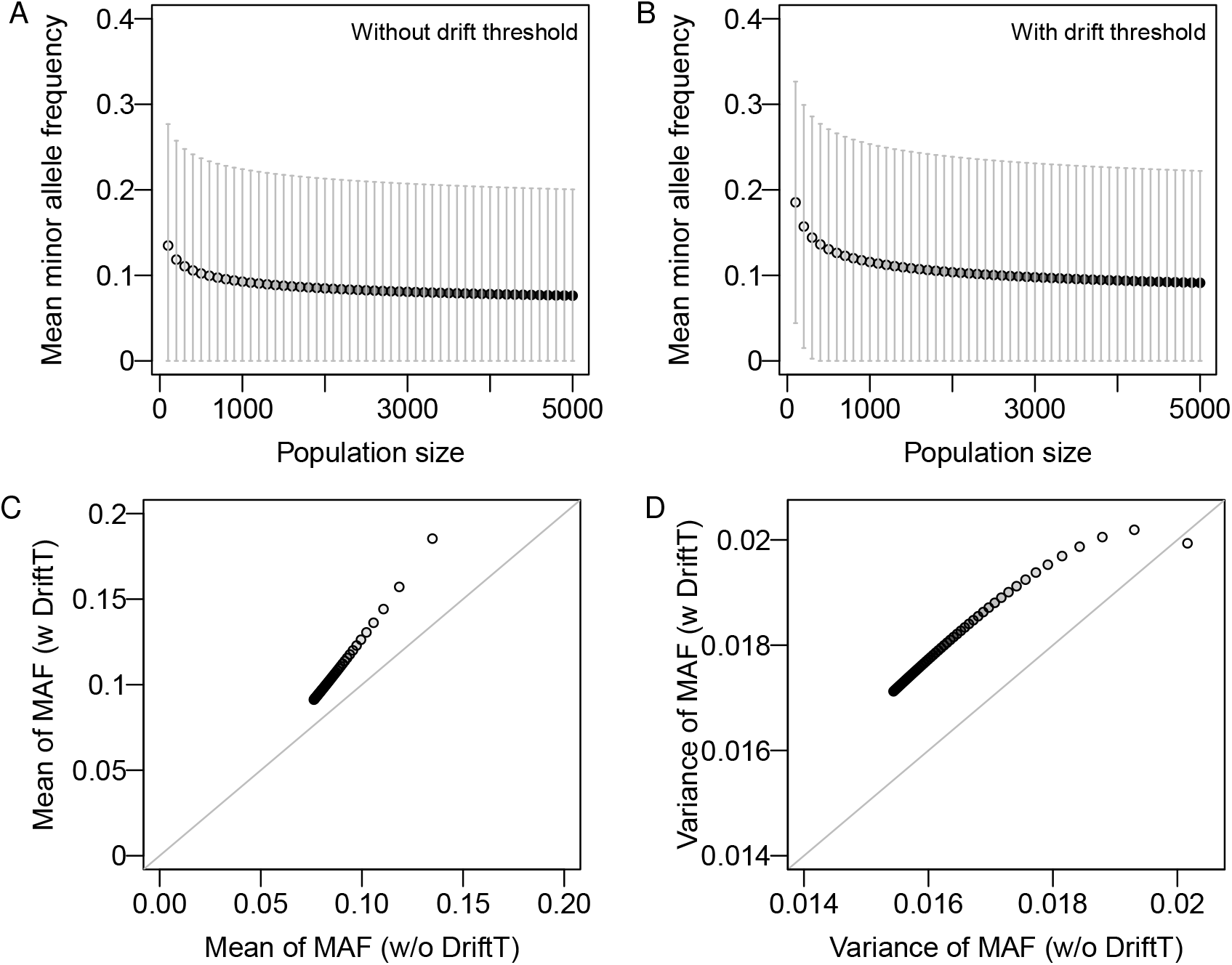
Mean and variance of minor allele frequencies (MAF) in the initial population. Mean and standard deviation (SD, the square root of variance) of minor allele frequencies (MAF) are calculated from the folded neutral SFS (see Methods; Hudson 2015). The mean and one SD of MAF are plotted by population size (A&B). The SFS spans all possible allele frequencies (without drift threshold, A) or is truncated using the genetic drift threshold (*MAF* ≥3, DriftT). Smaller population sizes show larger means and wider SD. The mean (C) and variance (D) are slightly elevated under the genetic drift threshold, where the population size ranges from 100 to 5000 with a step size of 100, indicated by the transparency of the black color.

**Figure S2.**
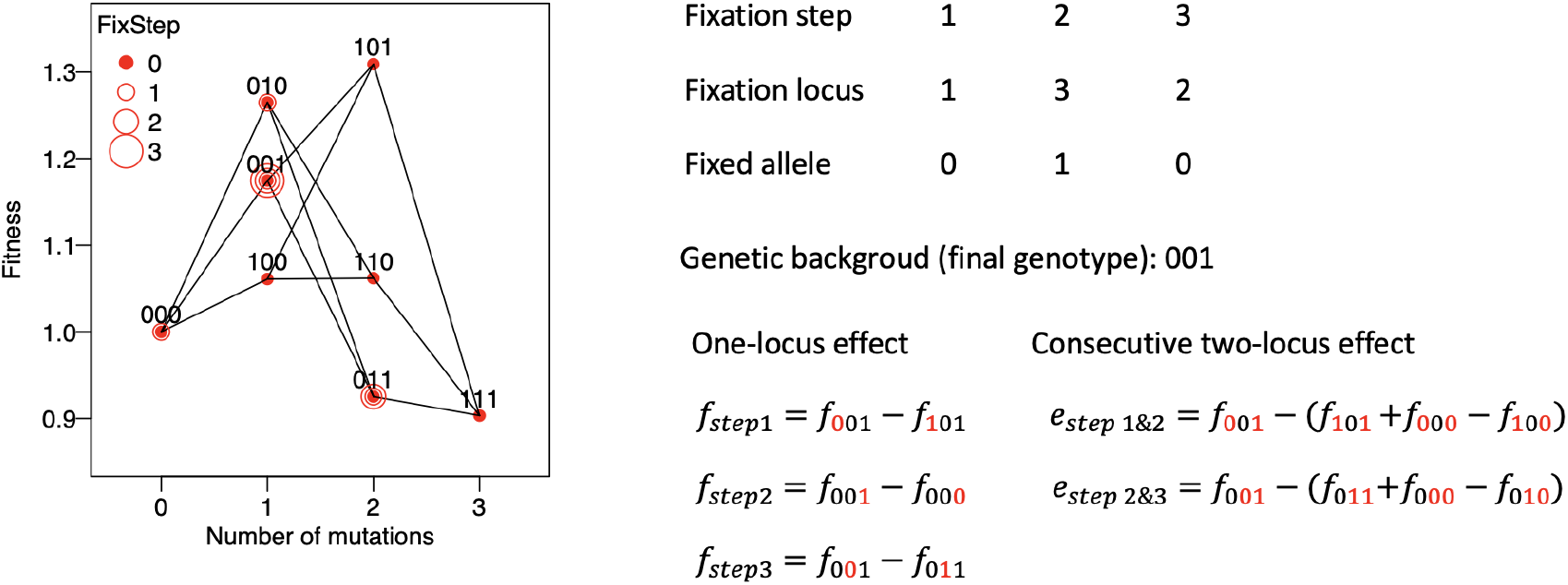
Example of selection coefficients along an adaptive path. We demonstrate in a three-locus fitness landscape how we calculate the one-locus effect and the pairwise epistasis (two-locus effect) along the adaptive path for a given reference genetic background. In this figure, the reference genetic background is the final genotype. The one-locus effect is the selection coefficient of the fixed allele compared to the lost allele (*e*.*g*., *f*_step 1_, *f*_step 2_, *f*_step 3_); the two-locus effect is the epistatic coefficient of the fixed genotype, composed of two consecutively fixed alleles (*e*.*g*., *e*_step 1 & 2_, *e*_step 2 & 3_). In this figure, each locus contains two alleles, 0 and 1. Genotype 000 is the reference genotype of the RMF model. The final genotype is 001. The alleles in red denote the fixed alleles at each step.

**Figure S3.**
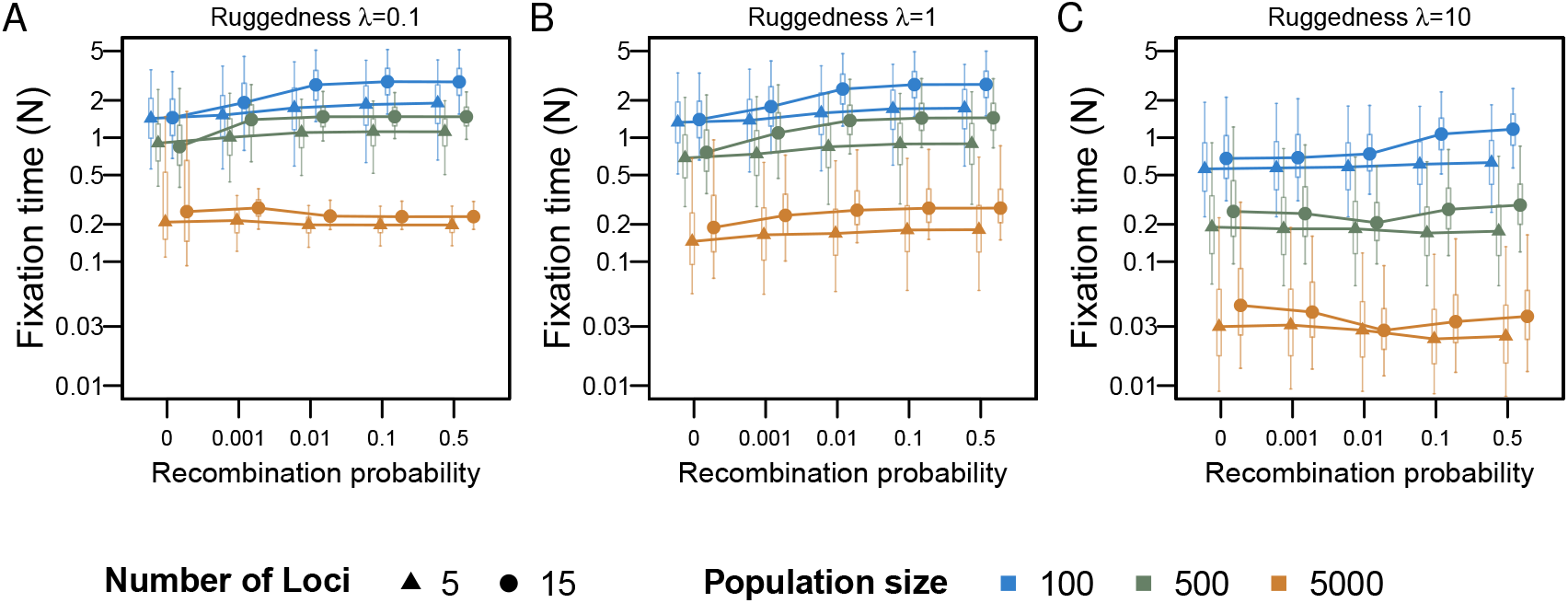
Distribution of fixation times scaled by the population size *N*. We consider two sizes of the fitness landscapes, 5 loci (*L* = 5, triangles) and 15 loci (*L* = 15, solid circles). The three population sizes (PopSize) are 100, 500 and 5000, which occupy a small or large fraction of genotype space (see Figure 2). Whereas there is no effect of recombination on fixation time on small fitness landscapes, the fixation time is variable across recombination probabilities on large fitness landscapes. The additive effect in the underlying RMF fitness landscape is *a* = 0.01. We study fitness landscapes of different ruggedness (0.1, 1, 10) by choosing three standard deviations of epistasis *σ* of 0.001, 0.01, and 0.1. Boxes represent the interquartile range, and whiskers extend outwards to cover 95% of the data.

**Figure S4.**
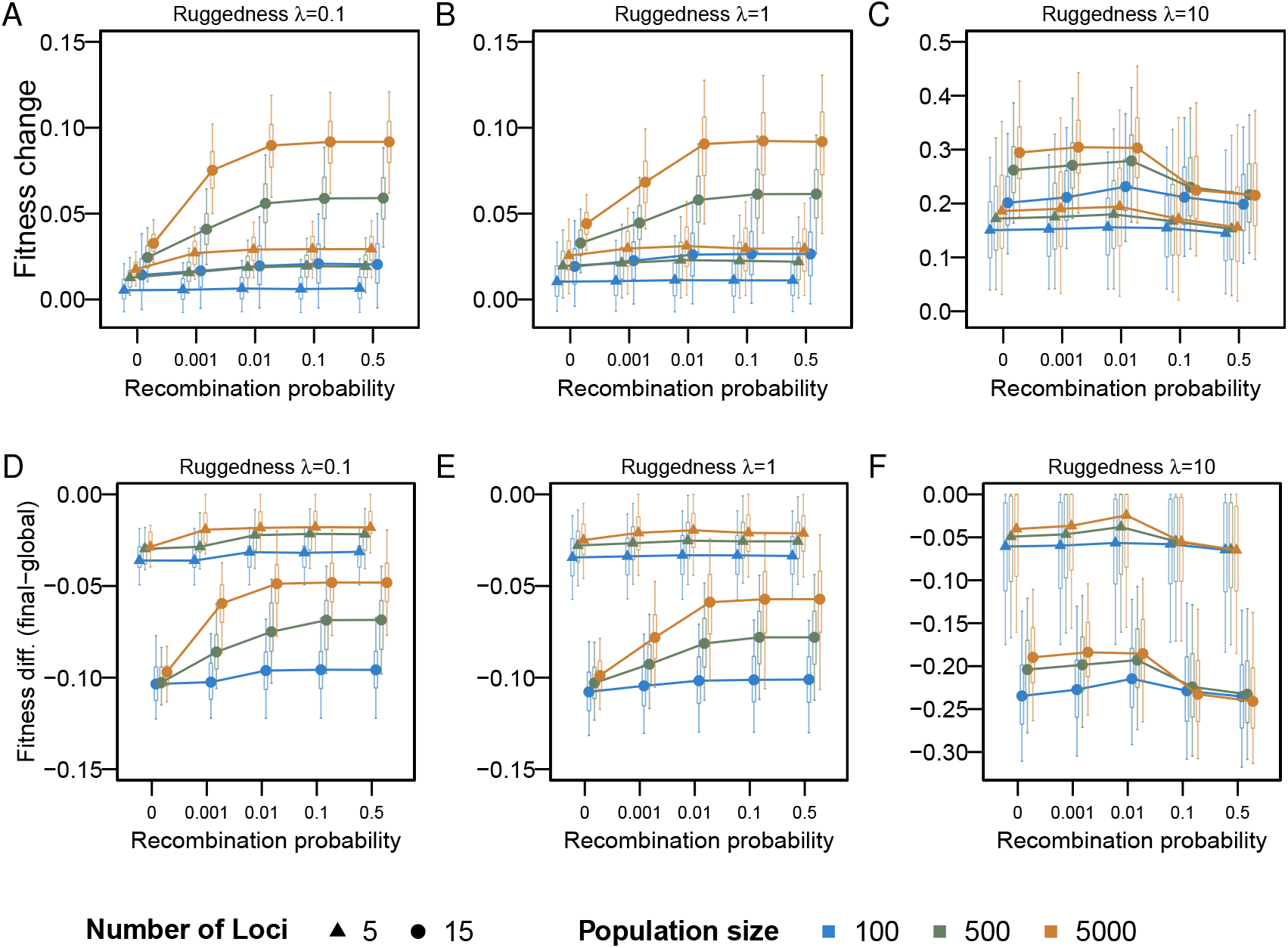
Fitness gain. The relative fitness change between the initial populations and the final genotype is defined as the absolute fitness change divided by the mean fitness of the initial population (A-C). D-E. Fitness difference between the final genotype and global peak (Fitness diff. (final-global)). Here, the fitness difference is defined as the fitness difference between the fixed genotype and the global peak, normalized by the fitness of the global peak. Boxes represent the interquartile range, and whiskers extend outwards to cover 95% of the data. Simulation parameters are as in Methods and Figure S3.

**Figure S5.**
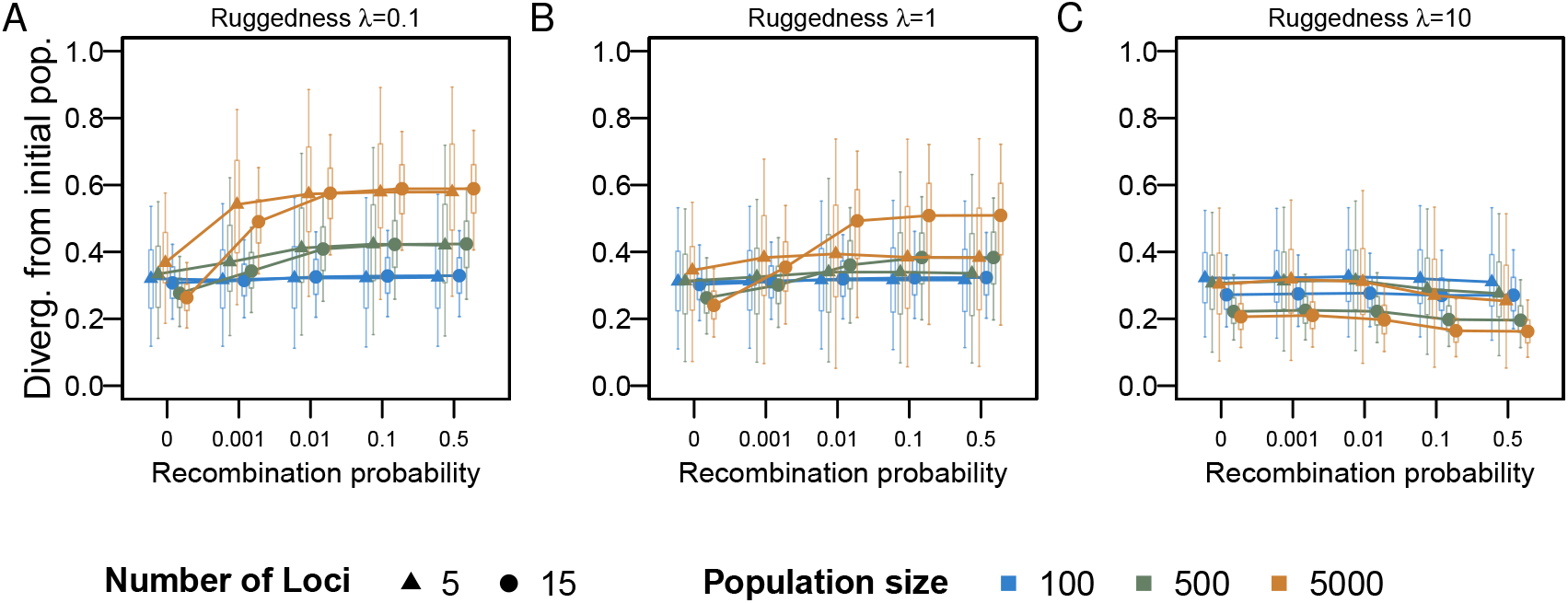
Distribution of the normalized Hamming distances between the final genotype and the initial population. Boxes represent the interquartile range, and whiskers extend outwards to cover 95% of the data. Simulation parameters are as in Methods and Figure S3.

**Figure S6.**
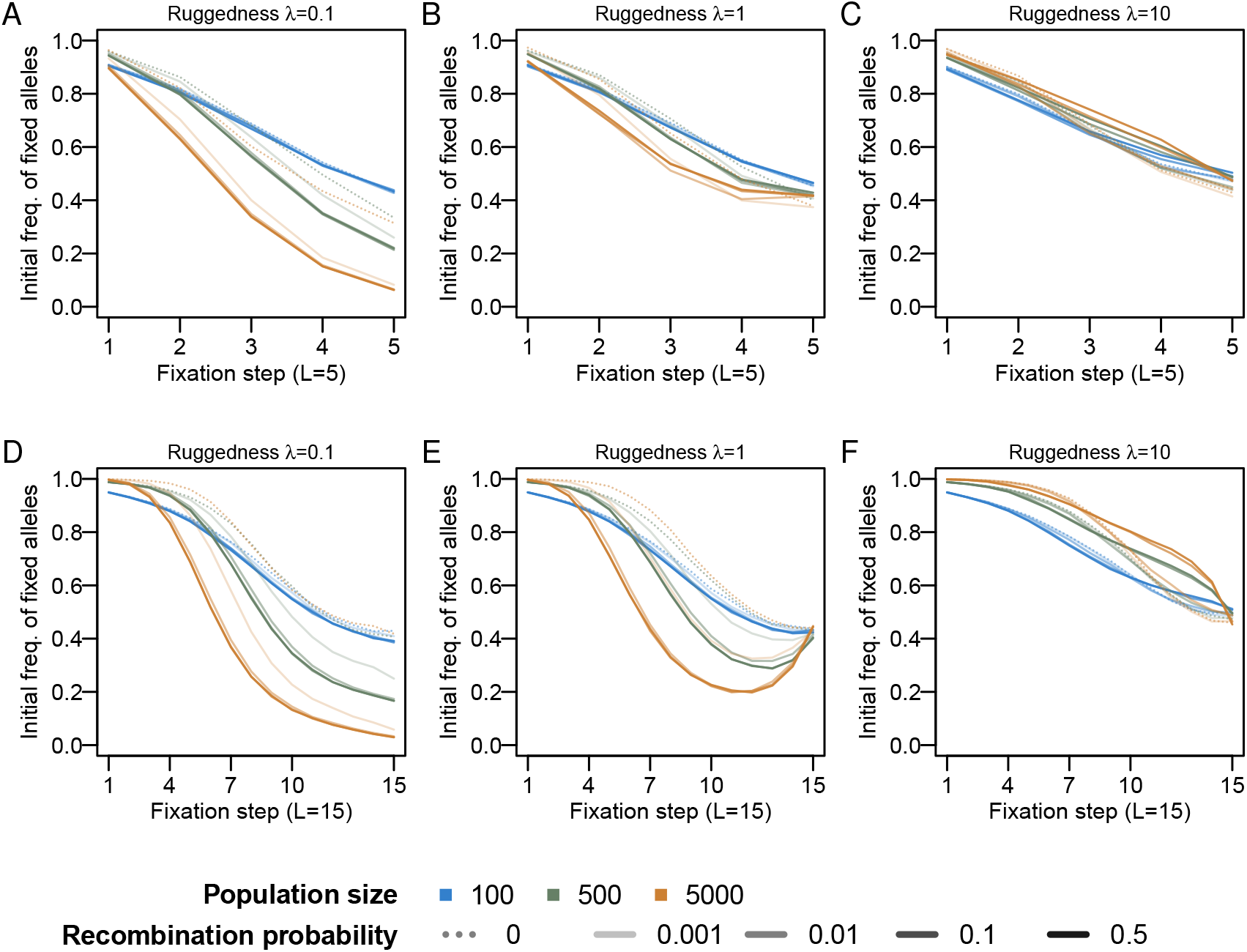
Mean of the initial frequency of fixed alleles on the adaptive path. The fitness landscape contains 5 loci (A-C) and 15 loci (D-F). The dotted line indicates no recombination. Simulation parameters are as in Methods and Figure S3.

**Figure S7.**
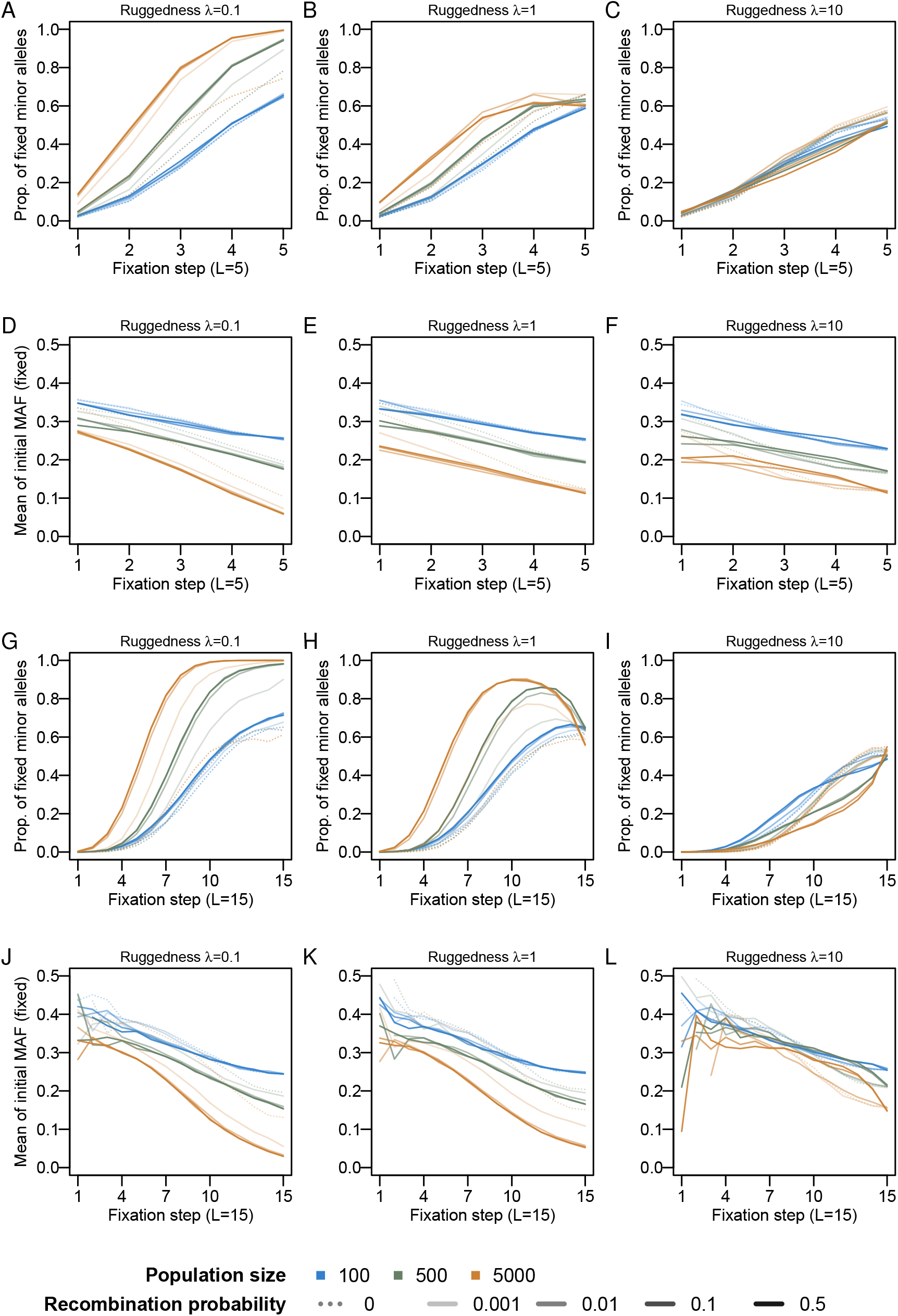
Characterization of fixed minor alleles. The fitness landscape contains 5 (A-F) or 15 loci (G-L). A-C & G-I Mean fraction of fixed minor alleles. D-F & J-L Mean of initial allele frequencies for fixed minor alleles. Fewer minor alleles are fixed at higher ruggedness, and the minor alleles that fixed tend to have higher initial frequencies. The dotted line indicates no recombination. Simulation parameters are as in Methods and Figure S3.

**Figure S8.**
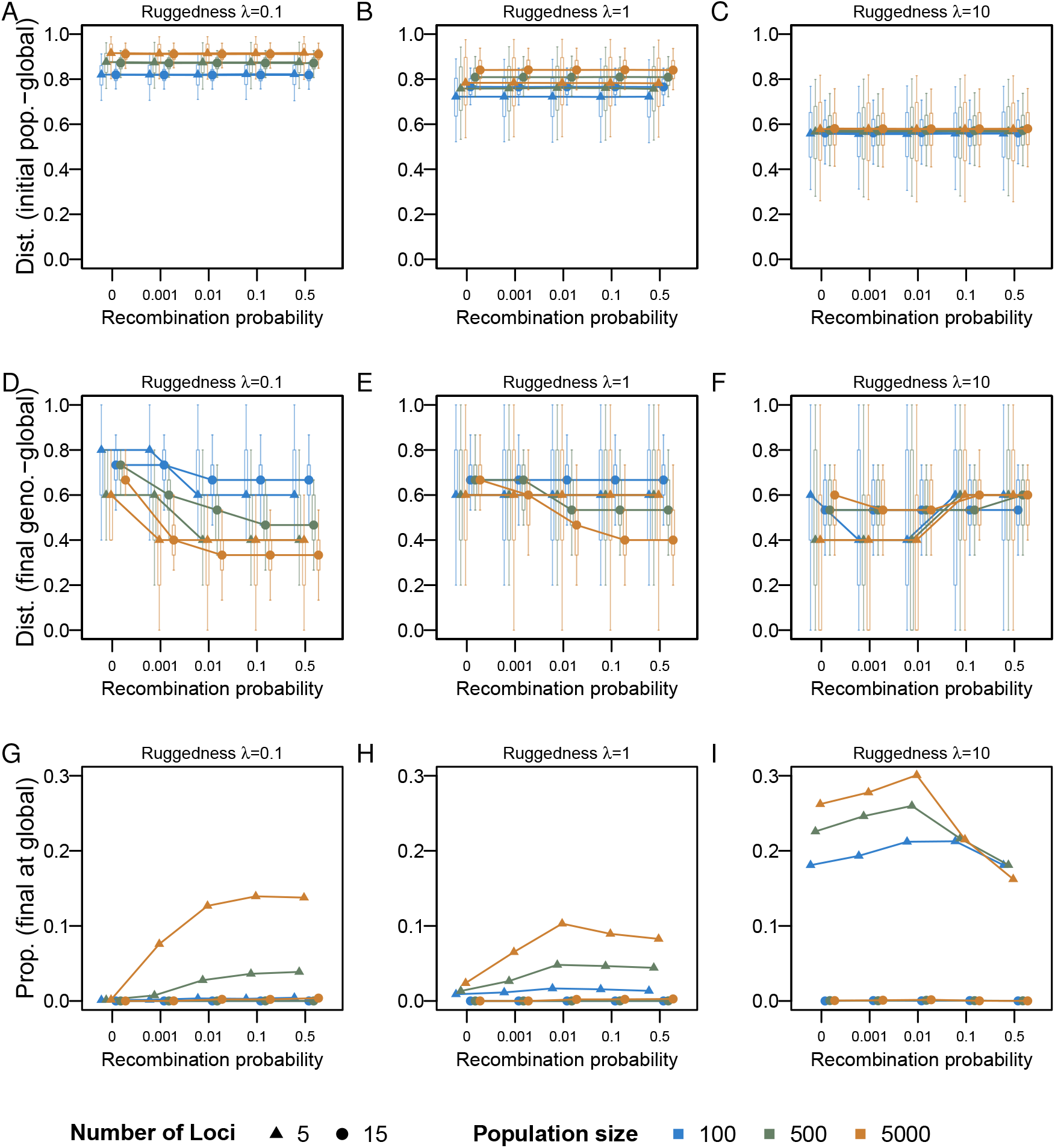
Genetic distance to global peak. A-C. Hamming distance between the initial population and global peak. The initial population is closer to the global peak at high ruggedness. D-F Hamming distance between the final genotype and global peak. High recombination only facilitates the population closer to the global peak at low ruggedness. G-I. Proportion of final genotype as the global peak. A population can achieve the global peak on a small fitness landscape. All the plots show the feature (y-axis) agaist recombination probability (x-axis). Boxes represent the interquartile range, and whiskers extend outwards to cover 95% of the data. Simulation parameters are as in Methods and Figure S3.

**Figure S9.**
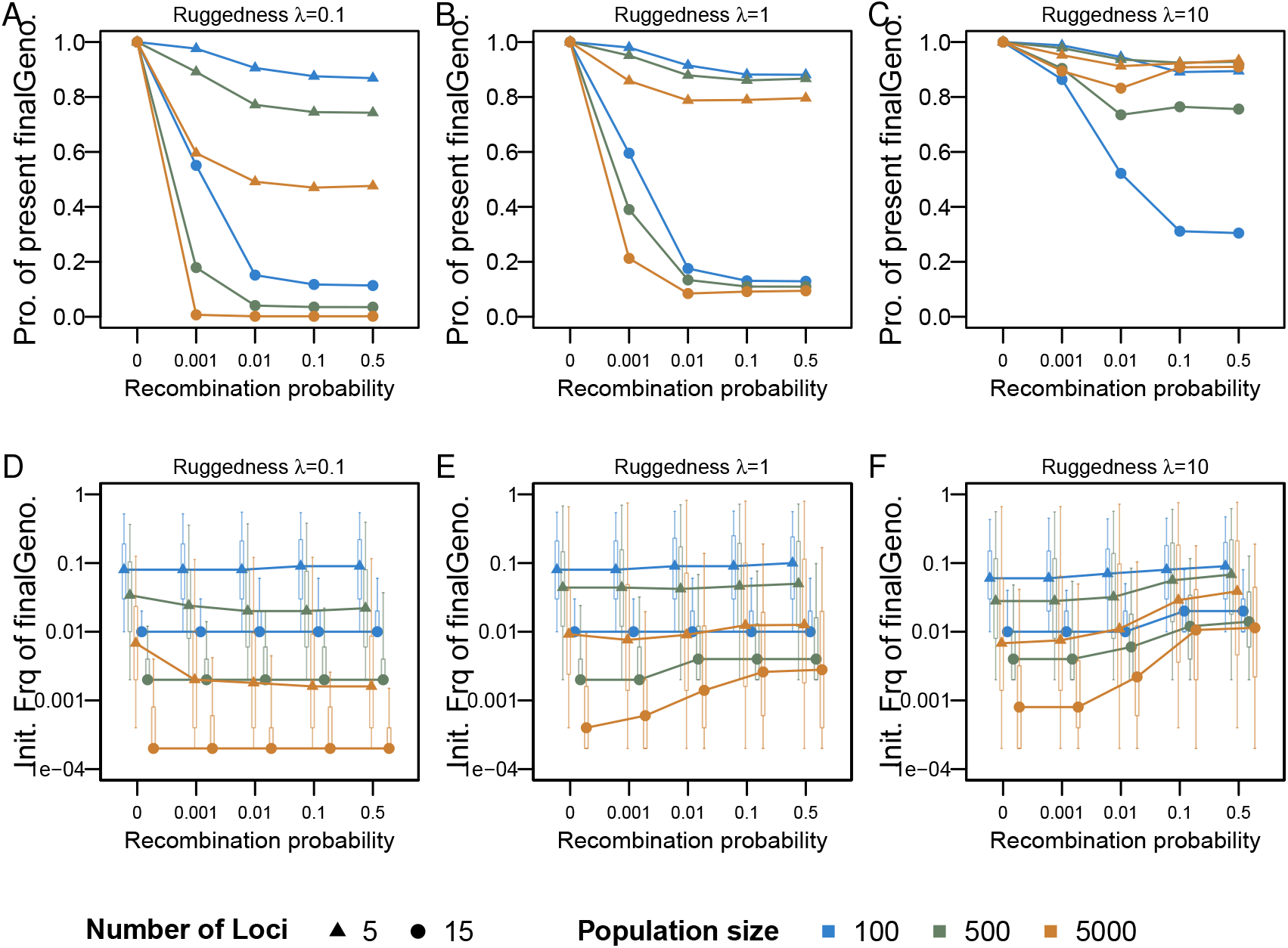
Presence of the final genotype in the initial population. A-C. Proportion of simulations in which the final genotype is present in the initial population. D-F. Distribution of initial frequencies of the final genotype. Here, only genotypes with at least one copy in the initial population are included. Boxes represent the interquartile range, and whiskers extend outwards to cover 95% of the data. Simulation parameters are as in Methods and Figure S3.

**Figure S10.**
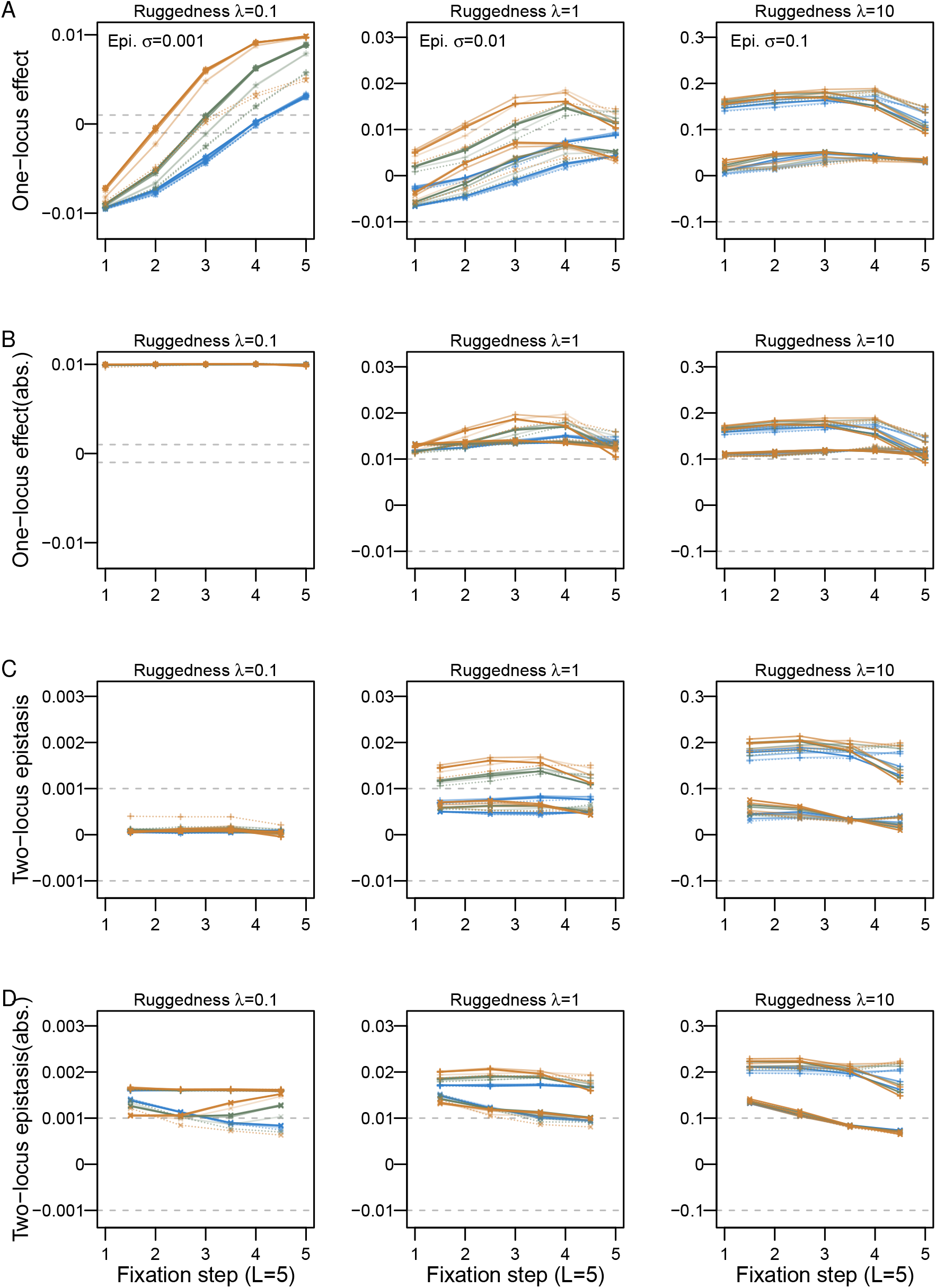
Selection coefficients along the adaptive path. The fitness landscape contains 5 loci. The mean of direct selection coefficients (one-locus effect, A) and two-locus epistasis (C) at each step is obtained from the genetic background using the major (crosses) or final (pluses) genotype. The means of their absolute values are plotted in B&D. Simulation parameters are as in Methods, Figure 5 and S3.

**Figure S11.**
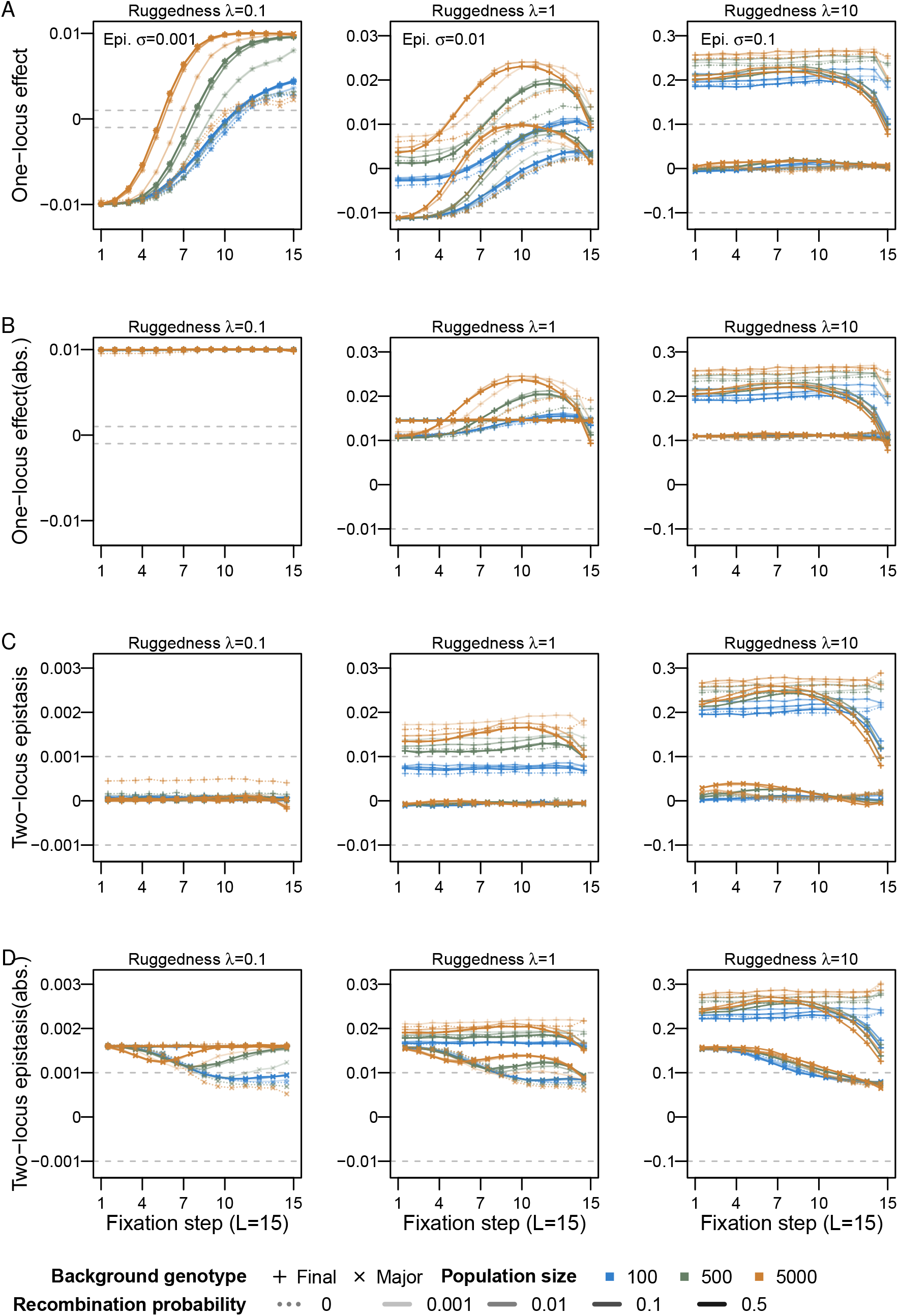
Selection coefficients along the adaptive path. The fitness landscape contains 15 loci. The mean of direct selection coefficients (one-locus effect, A) and two-locus epistasis (C) at each step is obtained from the genetic background using the major (crosses) or final (pluses) genotype. The means of their absolute values are plotted in B&D. Simulation parameters are as in Methods, Figure 5 and S3.

**Figure S12.**
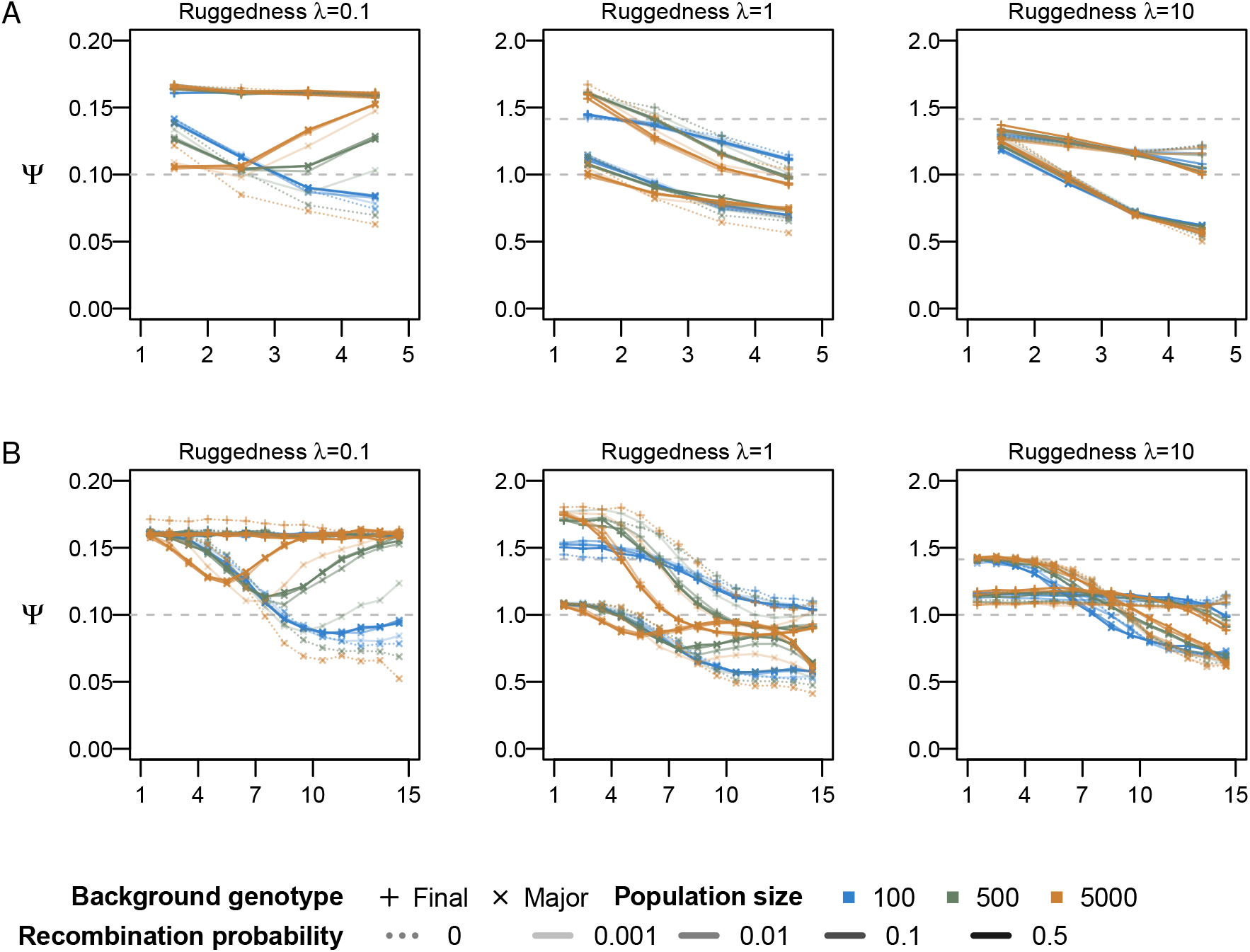
The ratio (Ψ) of means of absolute pairwise epistasis and absolute one-locus effects along the adaptive path. For detailed description, see Figure 5.

